# River network connectivity and postglacial history shape pollen- and seed-mediated gene flow across riparian populations of *Myricaria germanica*

**DOI:** 10.64898/2026.01.14.699434

**Authors:** Tania Chavarria-Pizarro, Christoph Scheidegger, Kailin Sun, Silke Werth

## Abstract

River network connectivity and postglacial history jointly shape patterns of pollen-and seed-mediated gene flow in riparian plants, yet their relative contributions across spatial and temporal scales remain incompletely understood. *Myricaria germanica* Desv., a pioneer riparian shrub formerly widespread along European rivers but now restricted to fragmented headwaters, provides an ideal model to investigate how river systems structure genetic connectivity. We analysed genetic diversity, population structure, and migration across 2,212 individuals from 67 populations spanning 12 Central European river catchments using 20 nuclear and six chloroplast microsatellite loci. By integrating biparentally inherited nuclear markers with maternally inherited chloroplast markers, we disentangled pollen- and seed-mediated gene flow across historical and contemporary timescales. Both marker systems revealed low genetic diversity, high inbreeding, and strong population differentiation. Nuclear microsatellites showed significant isolation by distance and extensive historical connectivity, with coalescent analyses indicating high pollen-mediated gene flow among catchments and identifying the Rhine and Danube as major long-term sources of migrants. In contrast, chloroplast microsatellites exhibited stronger spatial structure, limited admixture, and highly directional historical seed dispersal, consistent with constrained hydrochorous dispersal routes. Contemporary migration analyses further showed that present-day seed-mediated gene flow is largely confined within catchments, despite widespread historical pollen connectivity. Together, these results support a two-phase postglacial history in *M. germanica*, involving rare, directional seed dispersal during recolonization followed by prolonged pollen-mediated gene flow. Our findings highlight catchments as biologically meaningful management units and underscore the importance of conserving river network connectivity to preserve both the evolutionary legacy and long-term adaptive potential of riparian populations.

## 1 | INTRODUCTION

*Myricaria germanica* Desv. is a pioneer riparian shrub. specializing in open habitats and gravel soils close to unstable water regimes with sufficient water supply for seed germination (Werth et al. 2014; Sitzia et al. 2021). Furthermore. *M. germanica* is an indicator species of such gravel soil rivers. which are usually shaped by natural disturbances (Sitzia et al. 2021). This species used to be widespread along the upper and middle reaches of European rivers but is now a rare species confined to the headwaters of rivers which have not been modified by humans (Werth et al. 2014). Riparian plants like *M. germanica* depend on the linear flow of rivers to sustain their populations. Human activity has had a high impact on the linear flow of rivers, through building lateral embankments, channelizing the riverbeds with the purpose of irrigation, navigation, or to prevent land flooding (Jansson et al. 2000; Werth et al. 2014; Lind et al. 2019; Fink et al. 2022).

Dispersal in riparian plants is inherently asymmetric because pollen and seeds are transported by different vectors and operate over contrasting spatial and temporal scales. Pollen dispersal, often mediated by wind or insects, can connect populations across large geographic distances and across watershed boundaries, thereby reducing genetic differentiation in biparentally inherited nuclear genomes (Ennos 1994; Petit et al. 2005). In contrast, seed dispersal in riparian environments is frequently constrained by hydrological networks, occurring predominantly along river corridors via hydrochory and typically exhibiting strong directionality and episodic long-distance events (Nilsson et al. 2010; Werth et al. 2014; Auffret et al. 2015). As a consequence, maternally inherited chloroplast genomes often display stronger spatial genetic structure than nuclear markers, reflecting restricted seed movement and historical colonization pathways rather than contemporary pollen flow (Petit et al. 2005; Zhang & Hewitt 2003). Such decoupling between pollen- and seed-mediated gene flow has been documented in numerous riparian and woody plant species and provides a powerful framework for disentangling historical from contemporary connectivity in riverine landscapes (Ennos 1994; Ngeve et al. 2017). In species such as *M. germanica*, which combine selfing, hydrochorous seed dispersal, and the potential for long-distance pollen flow, contrasting genetic signals between nuclear and chloroplast markers are therefore expected and can yield critical insights into both postglacial recolonization dynamics and present-day dispersal constraints.

Assessment of contributions of historical climatic dynamics to population genetic structure could provide significant insights into diversification of species and their conservation (Hedrick 2005; Lee & Mitchell-Olds 2011; Meirmans 2012; Wang & Bradburd 2014; Tian et al. 2020). For example, climate oscillations in the quaternary occurred at dramatic speed (Dansgaard et al. 1993; Birks & Ammann 2000; Schönswetter et al. 2005), leading to extreme environmental changes, which caused substantial switches in the distribution of biota (van Andel & Tzedakis 1996). In Europe, the phylogeographic patterns suggest that lowland organisms survived in glacial refugia located along the southwestern, southern, eastern and northern border (Taberlet et al. 1998; Petit et al. 2003; Petit et al. 2005b; Hewitt 2004; Schönswetter et al. 2005). Simultaneously, mountain plants used glacial refugia present in central Alpine areas on ice-free mountain tops (Parisod 2022). *Myricaria germanica* has a wide distribution, from high altitude riparian habitats (e.g. glacier forelands) down to sea level (Sitzia et al. 2021), hence it could survive cold periods of the quaternary using some of the glacial refugia mentioned above, which could influence its contemporary genetic structure.

Microsatellite markers. also known as simple sequence repeats (SSR) are short, tandemly repeated sequences varying by the number of repeats. Microsatellites are very valuable molecular tools for the assessment of genetic diversity and for linkage mapping. They are also highly recommended for assessments of contemporary population genetic structure, as they have biparental inheritance, mutate fast, and are highly reproducible (Morgante & Olivieri 1993; Streiff et al. 1999; Jones & Ardren 2003; Patreze & Tsai 2010; Perdereau et al. 2014). In angiosperms, microsatellites can be found in both nuclear genomic DNA and in chloroplast DNA. These differ in their inheritance, as the former is biparental and the latter is maternal (Grivet et al. 2006). Chloroplast microsatellites are especially useful to determine ancient historical relationships, as they have lower evolutionary rates, are distributed throughout non-coding regions, and are highly conserved across genera (Provan et al. 1998; Streiff et al. 1999; Jones & Ardren 2003; Petit et al. 2005; Papi et al. 2012). The combined analysis of biparentally and maternally inherited microsatellite markers is important to fully understand the patterns of genetic diversity, structure, and differentiation in plant species (Djedid et al. 2022).

The aim of this study was to quantify genetic diversity, population structure, and the relative roles of pollen- and seed-mediated gene flow in *Myricaria germanica* across major Central European river catchments, integrating nuclear and chloroplast microsatellite markers to disentangle contemporary versus historical processes.

Specifically, we aimed to (i) quantify genetic diversity and inbreeding across populations and catchments; (ii) characterize population genetic structure using nuclear and chloroplast microsatellites; (iii) compare pollen- and seed-mediated gene flow across historical and contemporary timescales.

## 2 | METHODS

### 2.1 | Sampling

Our sampling included 2212 individuals collected from 67 populations situated in 12 catchments of rivers in Central and Eastern Europe (Table 1, Fig 1). The sampling strategy followed Werth and Scheidegger (2014), and a part of the populations included in this previous study were reanalysed for the present study. The number of sampling sites reflected the population number of the species in each catchment. For sampling, we used apical tips of branches without flowers, to avoid including seeds or pollen deposited on the plant. Samples were stored on silica gel at room temperature until DNA extraction. In large populations, tissue was sampled from 40 adult plants along a transect through the population following the flowing direction of the river, keeping at least 2m between adjacent plants. In small populations, tissue was collected from all plants. For each population, GPS coordinates were recorded, and the population size was estimated. In small populations, the plants were counted. In large populations, the individuals occupying a part of the area were counted and this number was used for extrapolation to the total area; those latter estimates were provided as intervals.

**FIGURE 1.**
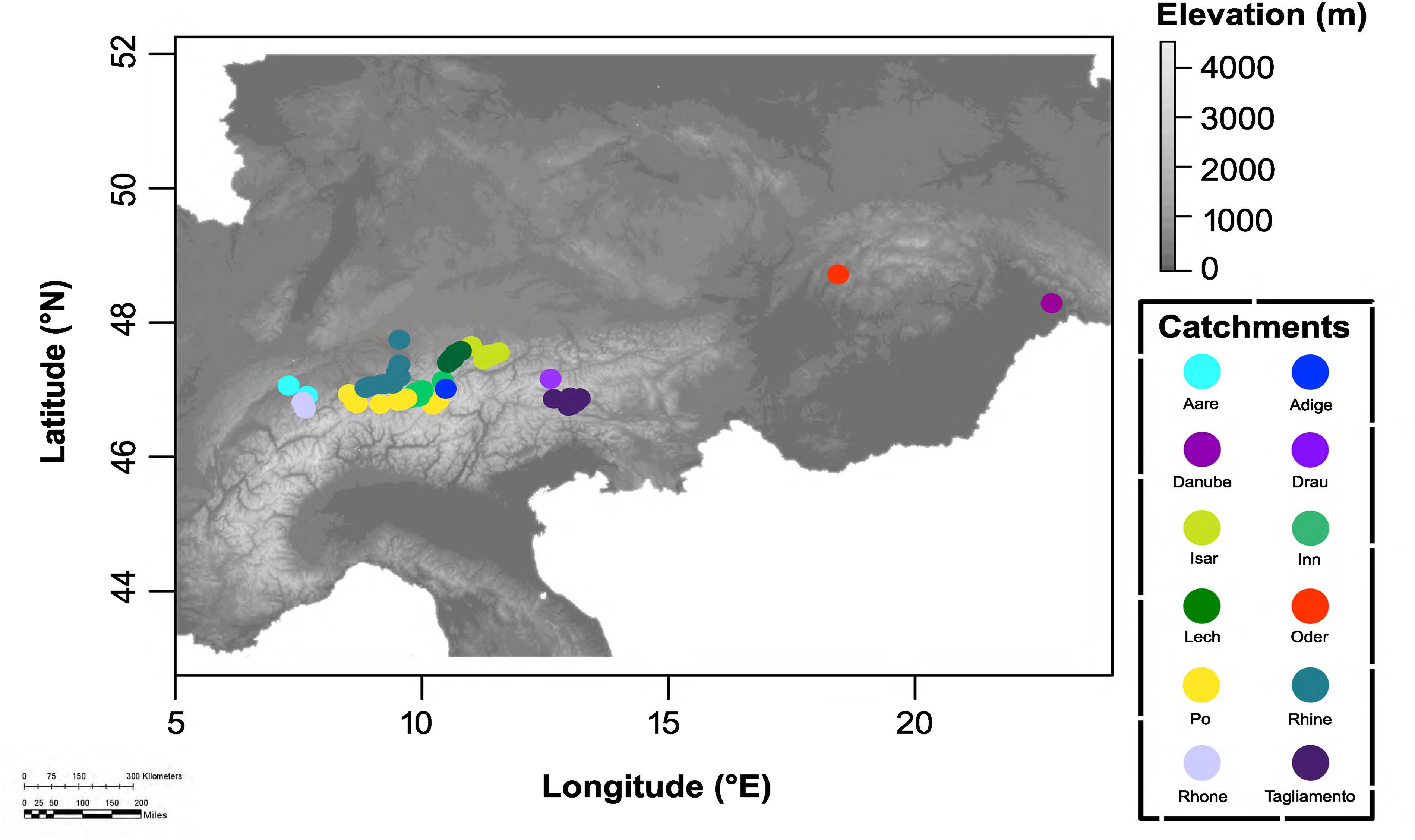
Map showing the geographic distribution of the 67 sampled populations of *Myricaria germanica* across 12 river catchments in Central Europe (2212 individuals). Populations are colour-coded according to their catchment of origin.

**Table 1.**
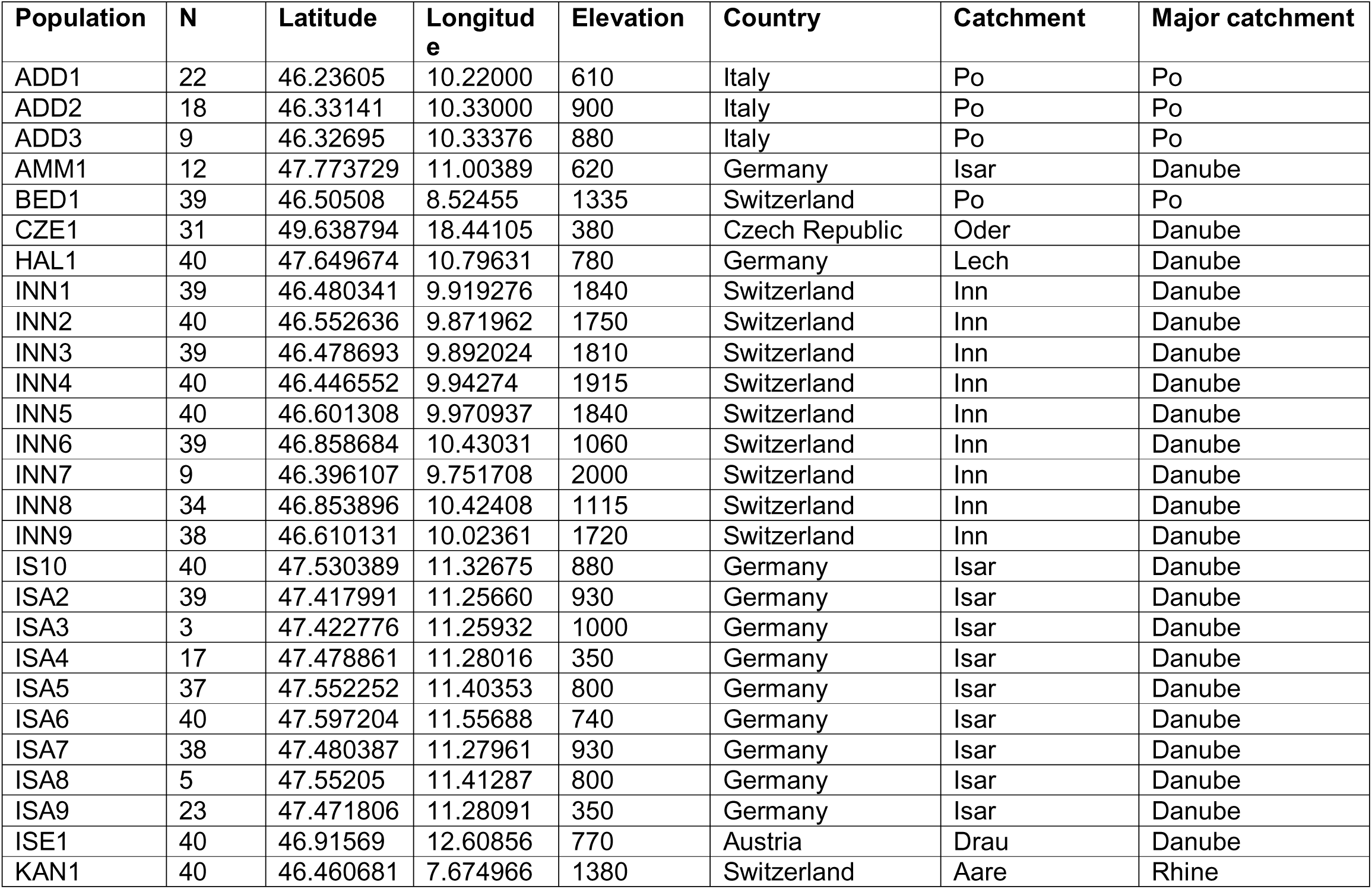
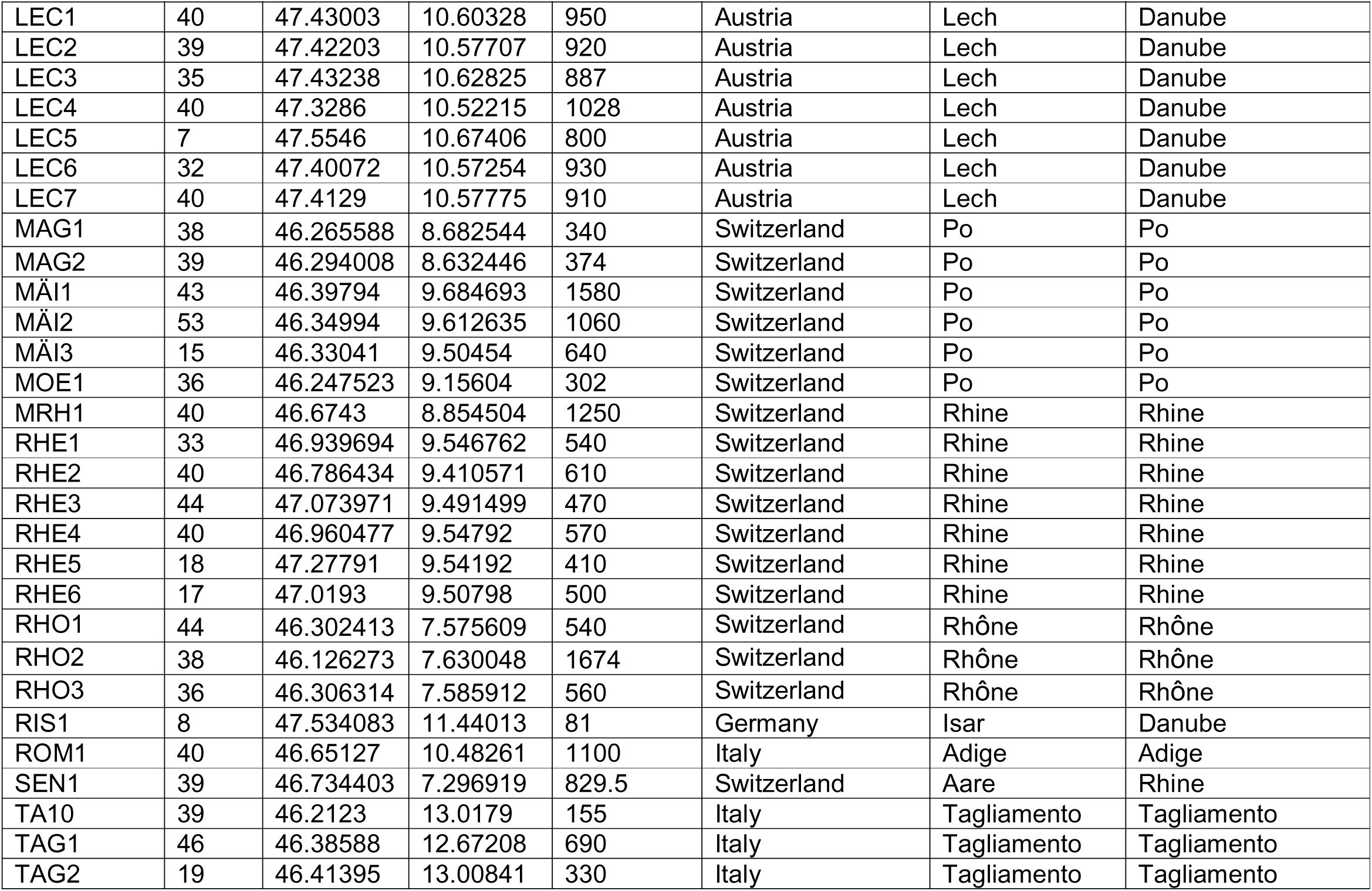
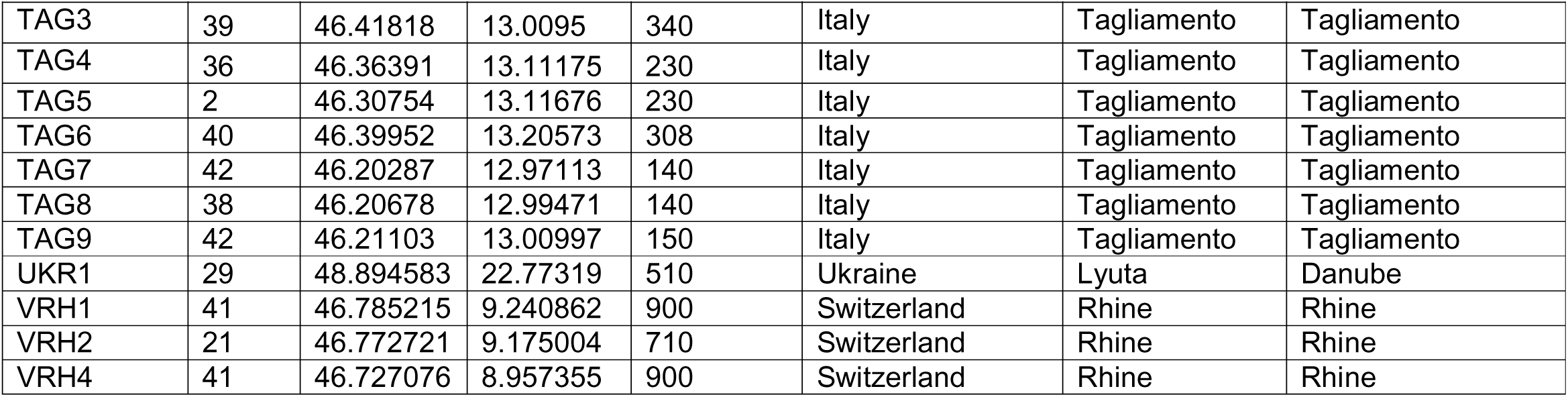
Location information for 67 populations of *M. germanica*. Shown are the number of samples per population (N). geographic coordinates (map datum WGS84). elevation (m). country. Catchment and Major catchment the populations belong.

Collecting permits were obtained from regional authorities when samples were collected in protected areas, but mostly we collected from public not protected areas.

### 2.2 | Genetic data and microsatellite methods

DNA was extracted with the DNeasy 96 plant kit (Qiagen). PCR, fragment analyses, and genotyping of 20 nuclear and six chloroplast microsatellites were performed according to the description in Werth and Scheidegger (2011) but leaving out the microsatellite loci Mg461 and Mg482 which proved difficult to amplify and had many missing genotypes.

### 2.3 | Genotypic Analysis

#### 2.3.1 | Levels of genetic diversity

As measures of genetic variability within populations, the mean number of alleles (Na), effective alleles number (Ne), the Shannon’s index (I), observed heterozygosity (Ho), and expected heterozygosity (He) were calculated using the R package ‘adegenet’ (adegenet_2.1.10) (Jombart & Ahmed 2011). Furthermore, Nei’s gene diversity (H), allelic richness (A_R_), and inbreeding coefficient (*F*_IS_) within populations were computed using the ‘Hierfstat’ R package (hierfstat_0.5-11) (Nei 1973. Goudet et al. 2002). For each locus, deviation from Hardy–Weinberg equilibrium (HWE) was tested, using Fisher’s exact tests implemented in adegenet (Jombart & Ahmed 2011). A heatmap was created to visualize HWE for each locus per population for nuclear data using R 4.5 (R Core Team 2019).

#### 2.3.2 | Genetic structure and population differentiation

The population genetic variance for both types of microsatellites (nuclear and chloroplast) was analyzed in GenoDive (version 3.0) (Meirmans 2020), which offers robust and flexible functional analysis of haploid datasets, treating them with the same precision and adaptability afforded to diploid and polyploid data. It supports seamless data import—including formats designed explicitly for haploid and dominant markers—and applies comprehensive statistical analyses across ploidy levels and efficient multithreaded processing make it especially well-suited for handling mitochondrial, chloroplast, or otherwise haploid genetic data (Meirmans 2020). With GENODIVE we calculated a global analysis of molecular variance (AMOVA), average F-statistics. pairwise fixation indices (*F*_ST_), and their significance levels (*p*-values) which were estimated with 1.000 permutations (Excoffier et al.2005. Weir & Cockerham 1984). Separate heatmaps were created to visualize all pairwise *F*_ST_ between the *M. germanica* populations for the nuclear and chloroplast microsatellites. respectively.

We tested for isolation by distance by regressing pairwise genetic distance [*F*_ST_ / (1 − *F*_ST_)] against the natural logarithm of geographical distance [km] across populations for the chloroplast and nuclear data and used a Mantel test (Mantel 1967) with 1.000 permutations implemented in the ‘adegenet’ package to determine if there was a correlation between genetic and geographic distance. Population structure was assessed using Bayesian model-based clustering implemented in STRUCTURE version 2.3.4 with admixed ancestry and the correlated allele frequency model (Pritchard. Stephens & Falush 2007). Twenty independent runs were performed for each K value, the number of genetic clusters, which ranged from 1 to 12 which where the number of catchments (the likelihood rapidly declined with increasing K values in preliminary runs after K=2) with a burn-in period of 50.000 iterations and 1.000.000 Markov Chain Monte Carlo (MCMC) replicates. The most likely number of clusters (K) was determined by considering the log-likelihood of each K value and using the ΔK method described by Evanno, Regnaut, and Goudet (2005). which was implemented in Structure Harvester (Earl & von Holdt 2012). The ΔK graph exhibits a maximum at the most likely value of K. These analyses allowed the detection of two distinct clusters with high admixture (see results). Because it is known that in the presence of a major grouping, STRUCTURE will not be able to detect finer structuring (Pritchard. Stephens & Falush 2007), we run a separate STRUCTURE analysis with each of the river catchments to identify subgroups. In these new analyses, we again considered K between 1 and 12.

Discriminant Analysis of Principal Components (DAPC) was used to identify subpopulations of the species, using the ‘adegenet’ package in R 4.5 (Jombart & Ahmed 2011. R Core Team 2019). The number of clusters was assessed using the function ‘find.clusters’, evaluating a range from 1 to 12. The optimal number of clusters was identified based on the Bayesian information criterion (BIC). as suggested by Jombart & Ahmed (2011). The function ‘find.clusters’ was used to transform original data into principal components (PCs). retaining 50 PCs in the analysis. The DAPC function performed a discriminant analysis using 10 PCs (>80% of variance explained), and ten eigenvalues were retained and examined. Ten independent runs were executed, and the average results were plotted.

We performed all the genetic structure analyses for all the populations (67) independently first. Then we grouped the populations by the catchment that they belonged to and performed the genetic structure analysis on those catchments. Our 67 *M. germanica* populations were found in 12 catchments (Fig 1, Table 1). In addition, we constructed phylogenetic trees for nuclear and chloroplast data using neighbor-joining (NJ) and Unweighted Pair Group Method with Arithmetic Mean (UPGMA) methods in R. with phangorn (Schliep 2011) and ape (Paradis et al. 2004). We used a bootstrap of 5000 using the function bootstrap.phyDat().

As we said above, the analysis of molecular variance was calculate using the software GENODIVE the same we used to estimate the F-statistics (Weir and Cockerham, 1984). AMOVA and associated significance levels (p-values) were estimated using 1,000 permutations (Excoffier et al. 2005; Weir & Cockerham 1984). Variance components obtained from the AMOVA were subsequently used to estimate the relative contributions of pollen- and seed-mediated gene flow to total gene flow, applying the approach described by Ennos (1994):

*Pollen flow/seed flow* =(1*ϕSC*(*B*) − 1) − 2 (1*ϕSC*(*M*) − 1|(1*ϕSC*(*M*) − 1)

Where ϕSC(B) and ϕSC(M) are levels of population differentiation calculated from biparentally (nuclear) and maternally (cpDNA) inherited markers, respectively. The calculation of the pollen–seed flow ratio presented here assumes that maternal inheritance of cpDNA is the rule in *M. germanica* as in most other angiosperms (Ennos et al. 1999).

#### 2.3.2 | Migration levels

To test specific models about the directionality of gene flow and to obtain Bayesian estimates of effective population sizes and bidirectional rates of migration in *M. germanica*, we used the coalescent-based software MIGRATE-n version 3.26 (Beerli & Felsenstein 2001; Beerli 2016) for nuclear data (diploid) and chloroplast data (haploid). MIGRATE-n makes the following assumptions: i) constant population size through time or random fluctuation around an average size; ii) individuals are randomly mating within populations; iii) the mutation rate is constant through time and is the same in all parts of the genealogy; iv) the immigration rate is constant through time; and v) the studied populations exhibit a recent divergence and not an old split, so they exchange material through gene flow. We ran MIGRATE-n to estimate migration rates among catchments (all parameters free to vary) with one long MCMC chain, sampling every 1000th step for a total of 200.000.000 genealogies after a burn-in of 200.000 steps in the chain. A Brownian motion model was used which approximates the stepwise mutation model commonly used for microsatellite data but converges faster than the standard stepwise mutation model. Populations were randomly resampled to 100 individuals to speed up convergence. Starting parameters for population size h and migration rates were inferred from *F*_ST_ values. mutation rate modifiers were deduced from the data using Watterson’s h. We performed a series of preliminary runs to explore run conditions and prior distributions for the data from each catchment. Bayesian estimation of migration rate and population sizes were run with one long chain and static heating (temperatures of four Markov chains: 1. 1.5. 3. 10.000.000). To facilitate comparison among migration pathways, we converted posterior median mutation-scaled migration rates (M) into relative migration probabilities scaled between 0 and 1. These probabilities represent the proportion of total inferred gene flow attributed to each migration route.

For every directed migration parameter *M_j_*→*_i_*:

Relative migration probability ∈ [0,1] is:

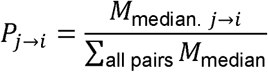

As we found that seed dispersal likely dominates over pollen in *M. germanica* contemporary migration patterns (see results), we calculate the recent migration rates between the 12 catchments with nuclear microsatellite data by implementing BA3-MSAT BayesAss Edition 3.0.5 (Wilson & Rannala 2003) and compared them with the relative migration rates obtain for the nuclear data with MIGRATE-n. First, we assessed the optimal mixing parameters for migration rates (deltaM = 0.1). allele frequencies (delta = 0.55). and inbreeding coefficients (deltaF = 0.0375) by running ten repetitions in BA3-MSAT- autotune V 3.0.5 as recommended by Mussmann et al. (2019). Subsequently. BA3-MSAT was run with the predefined mixing parameters for 50 million generations. sampling every 100th generation. The first million generations were discarded as burn-in and chain convergence was assessed in Tracer v.1.7.1 (Rambaut et al. 2018).

## 3 | RESULTS

### 3.1 | Genetic diversity

Across the 20 nuclear microsatellite loci. we detected a total of 169 alleles. with low to moderate null allele frequencies (0.023–0.17). As null allele frequencies were low. no loci were excluded from subsequent analyses. In total, 1.684 multilocus genotypes (MLGs) were identified. The number of alleles per locus ranged from three to 22, expected heterozygosity (Hs) ranged from 0.06 to 0.40, and F-statistics indicated substantial genetic structure (*F*_IS_ = 0.24–0.47; *F*_IT_ = 0.36–0.81; *F*_ST_ = 0.35–0.80; Table S1). At the population level the number of MLGs ranged from two to ten.

Shannon diversity indices ranged from 0 to 0.99, while Nei’s gene diversity varied between 0 and 0.50 (Table 2). Inbreeding coefficients (*F*_IS_) were high across populations (0.10–0.93). and population-level *F*_ST_ values ranged from 0.36 to 0.95 (Fig. S1A). Most populations showed high levels of homocigotes at multiple loci according to the Hardy–Weinberg equilibrium (p < 0.05; Fig. S2).

**Table 2.**
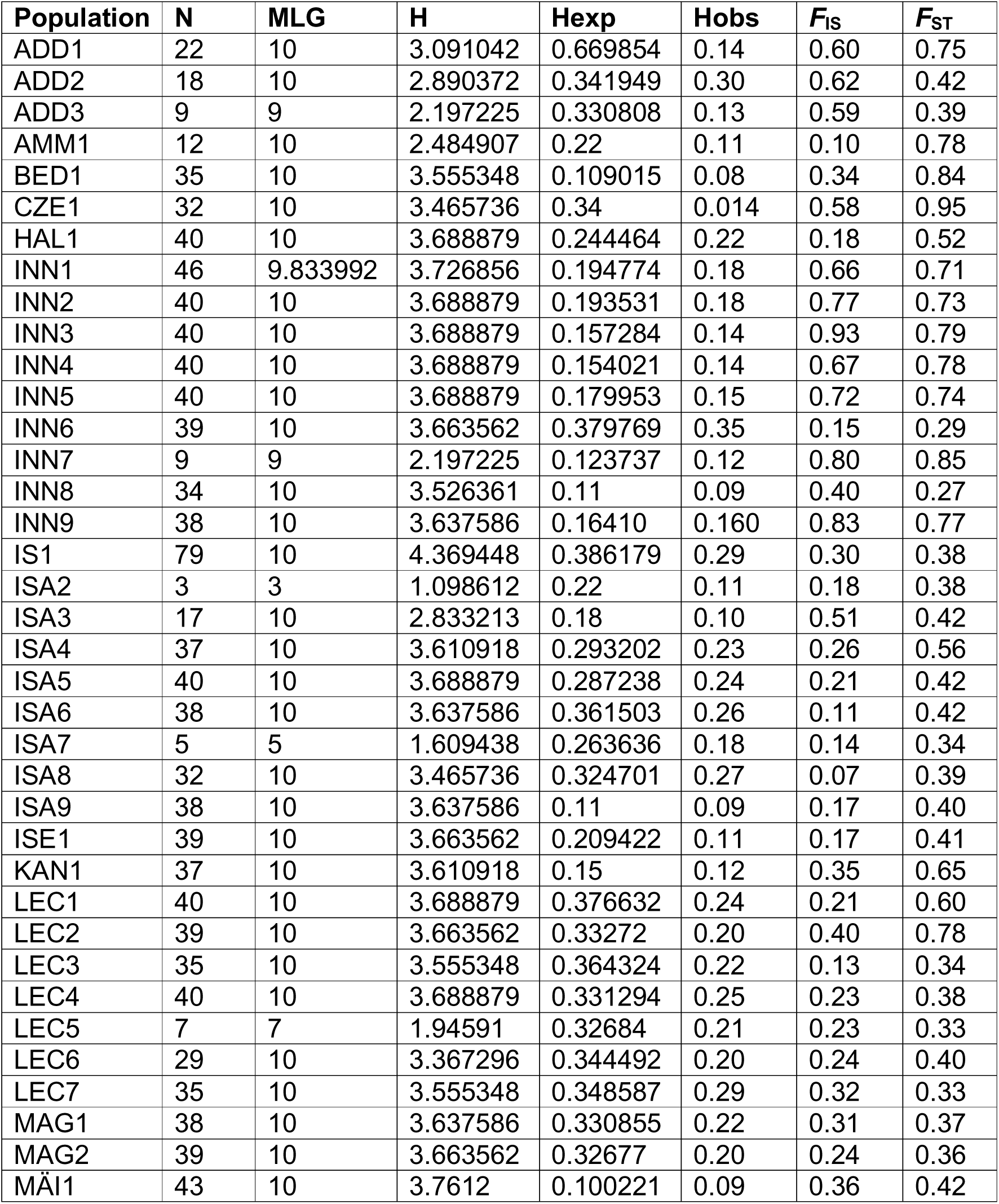
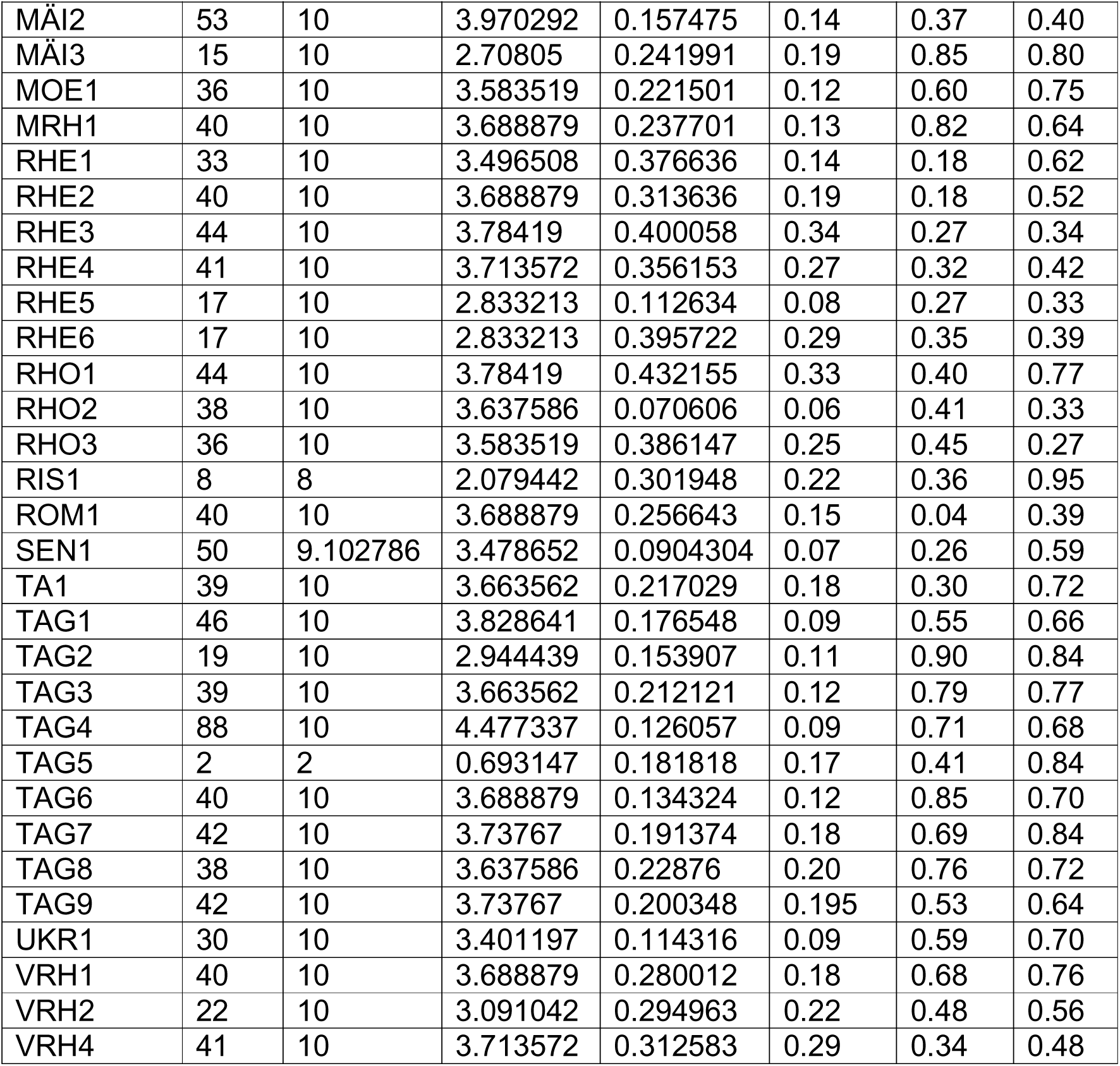
Population statistics for 67 geographically distinct populations of *M. germanica* based on twenty nuclear microsatellites. Shown are MLG. the number of multilocus genotypes; H. the Shannon-Wiener Diversity index; Hexp. expected heterozygosity; *F*_IS_. Wright’s inbreeding coefficient; *F*_ST_. fixation index of subpopulation to total.

Using six haploid chloroplast microsatellite loci, we detected 30 alleles and identified 55 chloroplast multilocus genotypes. The number of alleles per locus ranged from two to nine, and expected heterozygosity (Hs) ranged from 0.08 to 0.34. Estimates of genetic structure were high (*F*_ST_ = 0.10–0.95; Table S2). At the population level the number of chloroplast MLGs ranged from one to ten. Shannon diversity indices varied from 0 to 0.94, and Nei’s gene diversity ranged from 0.38 to 0.98 (Table 3).

**Table 3.**
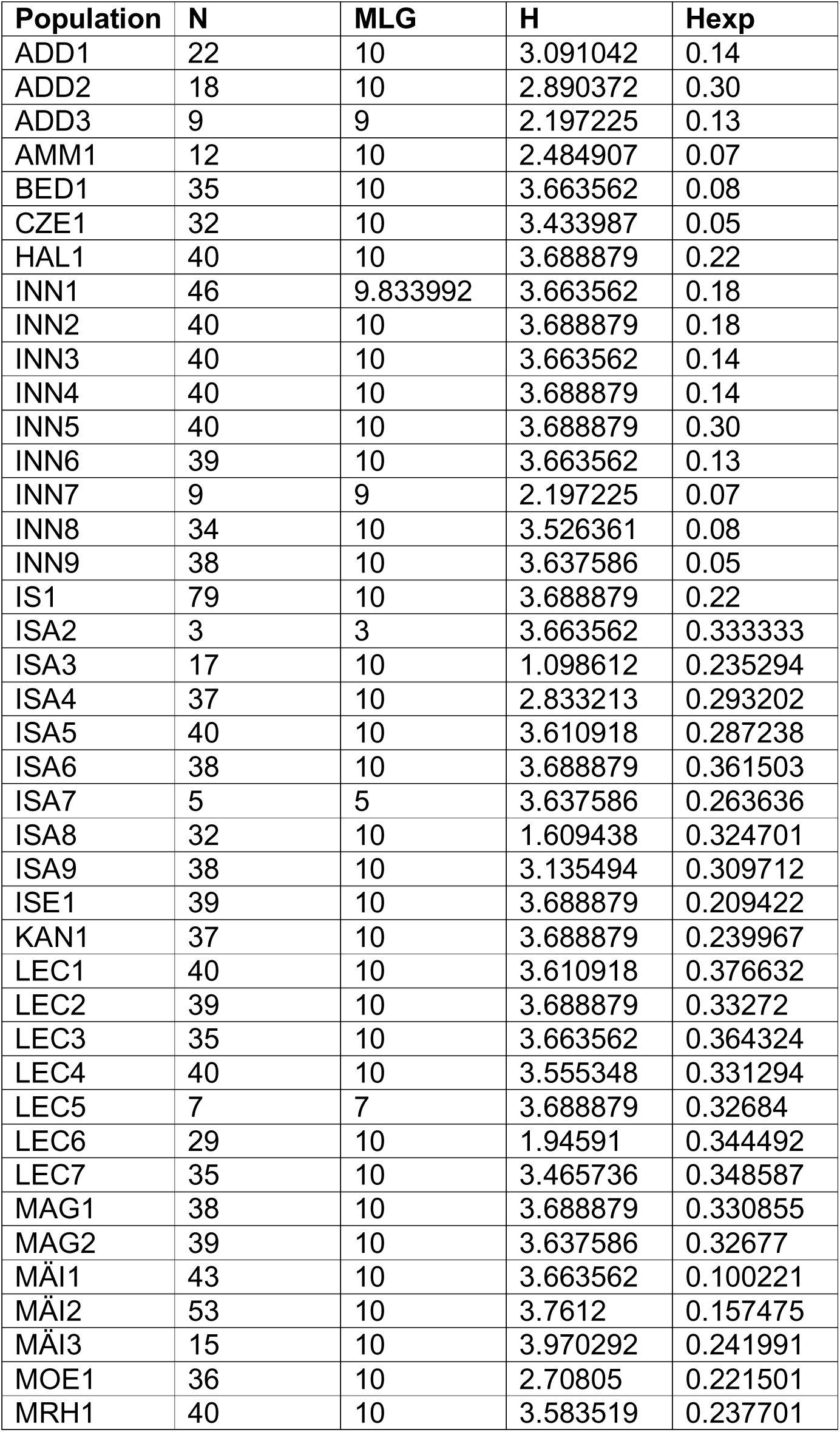
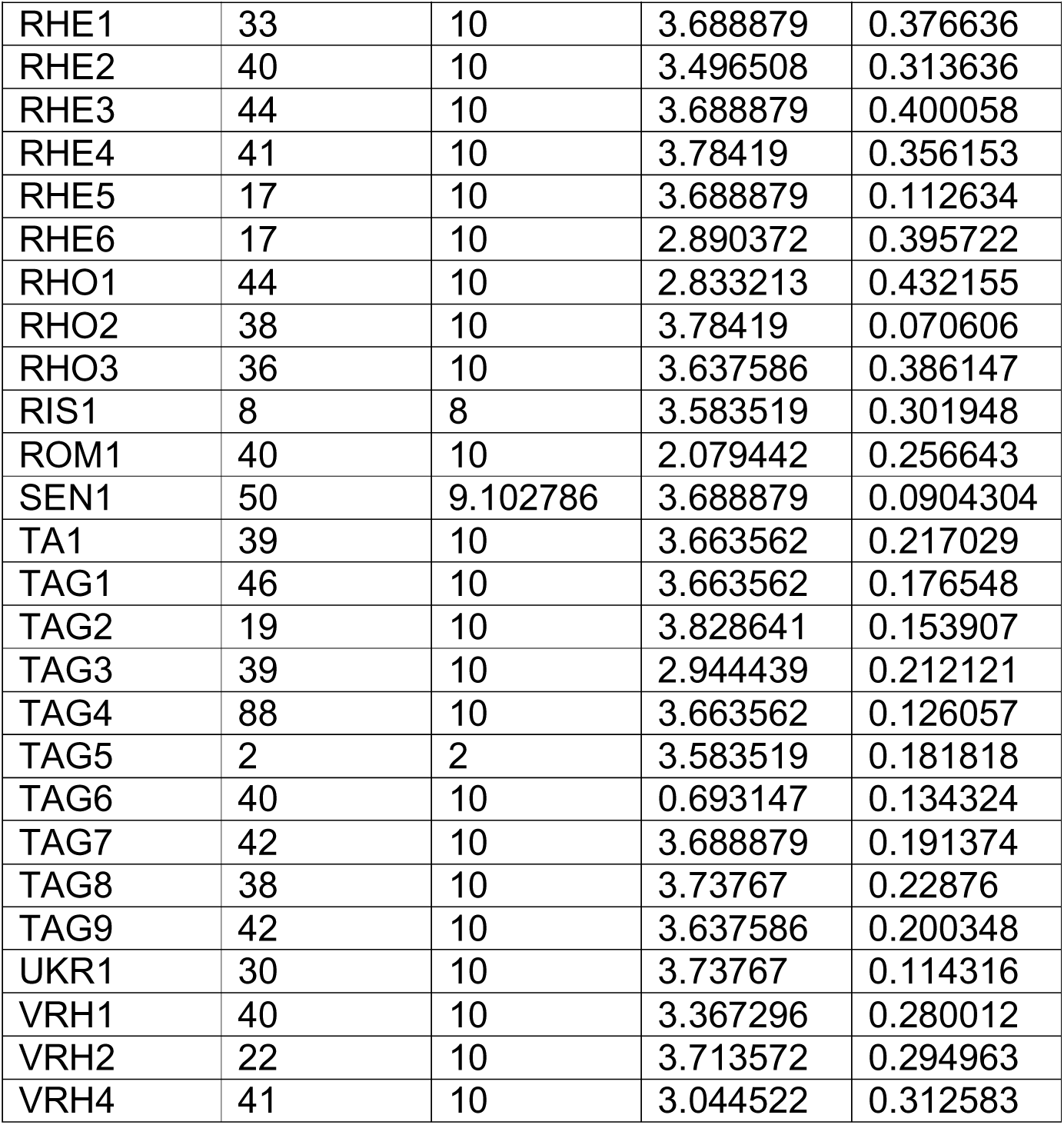
Genetic diversity and population differentiation for 67 populations of *M. germanica* based on haploid chloroplast microsatellites. Shown are sample size (N). number of haplotypes (Nh). Shannon–Wiener diversity index (H). haplotype diversity (Hd; Nei 1987). and population differentiation (FST).

**Table 4.**
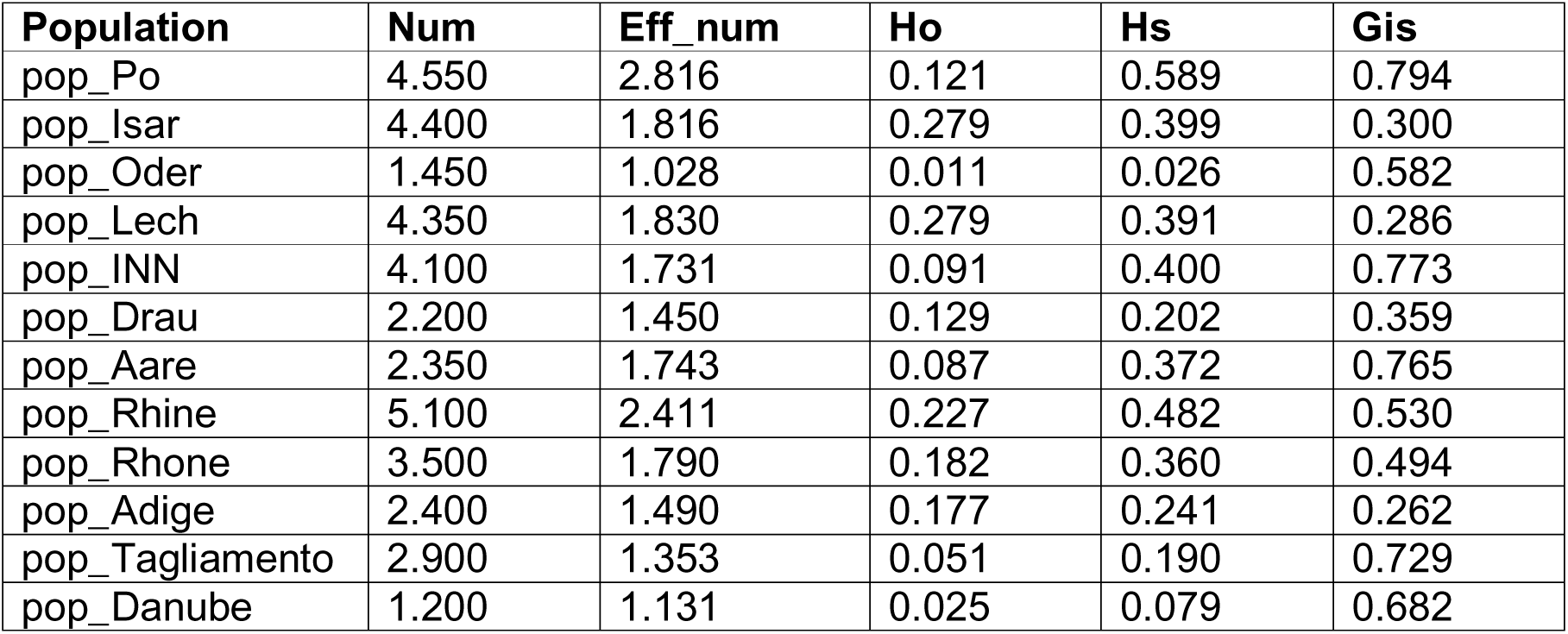
Indices of genetic diversity per catchment calculated with nuclear data (diploid). Num (number of alleles). Eff_num (effective number of alleles). Ho(Observe heterozigosity). Hs (expected heterozigosity). Gis (inbreeding coefficient).

**Table 5.**
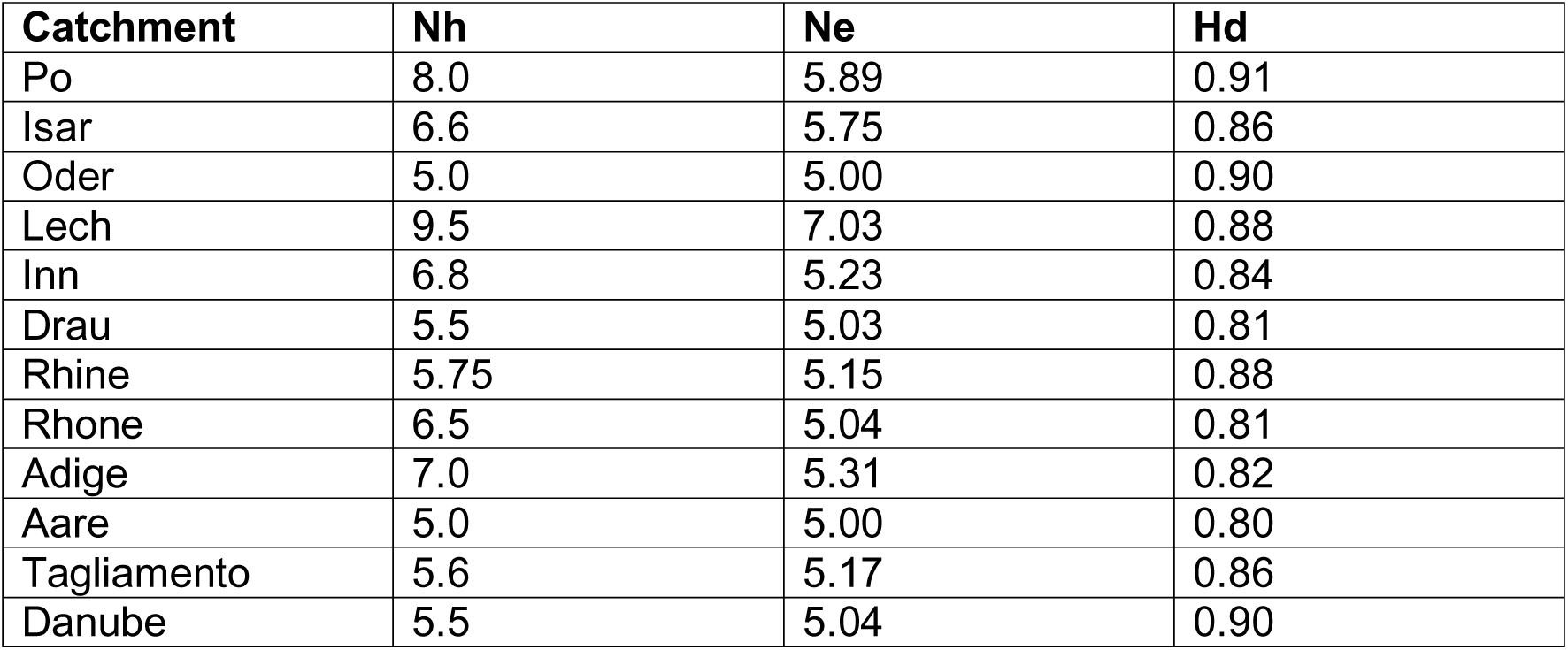
Genetic diversity indices per catchment based on haploid chloroplast microsatellites. Nh = number of haplotypes; Ne = effective number of haplotypes; Hd = haplotype diversity (Nei 1987).

Population-level *F*_IS_ ranged from –0.10 to 0.93. and *F*_ST_ ranged from 0.10 to 0.95. indicating strong differentiation among populations.

### 3.2 | Population genetic structure

#### 3.2.1 Nuclear microsatellites

Bayesian clustering analysis of nuclear microsatellites (STRUCTURE) identified two main genetic clusters (K = 2; Fig. 2A–B, Fig. 3). Most populations were dominated by a single cluster, although several showed varying degrees of admixture (Fig. 2A; Figs. S3–S15). Given the large dataset (67 populations, 2.212 individuals).

**FIGURE 2.**
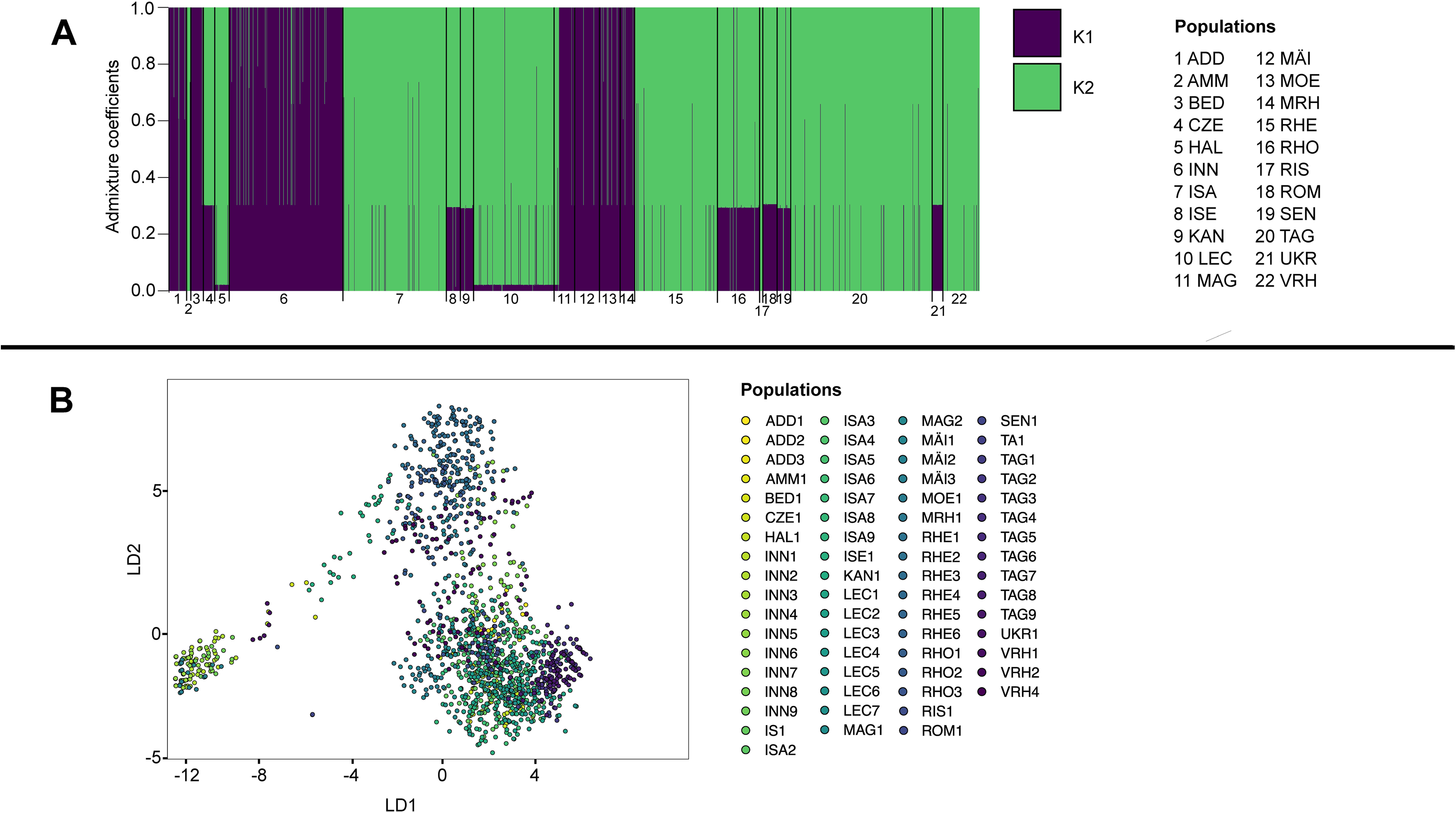
Population genetic structure of *M. germanica* based on 20 nuclear microsatellite loci. (A) Bayesian clustering analysis inferred with STRUCTURE. Each vertical bar represents an individual, and colours indicate membership coefficients for the two genetic clusters (K = 2) identified using the ΔK method. Individuals are grouped by population for visual clarity. (B) Discriminant analysis of principal components (DAPC) showing individual genotypes coloured by population. The first two discriminant axes explain 80% of the total discriminant variance.

**FIGURE 3.**
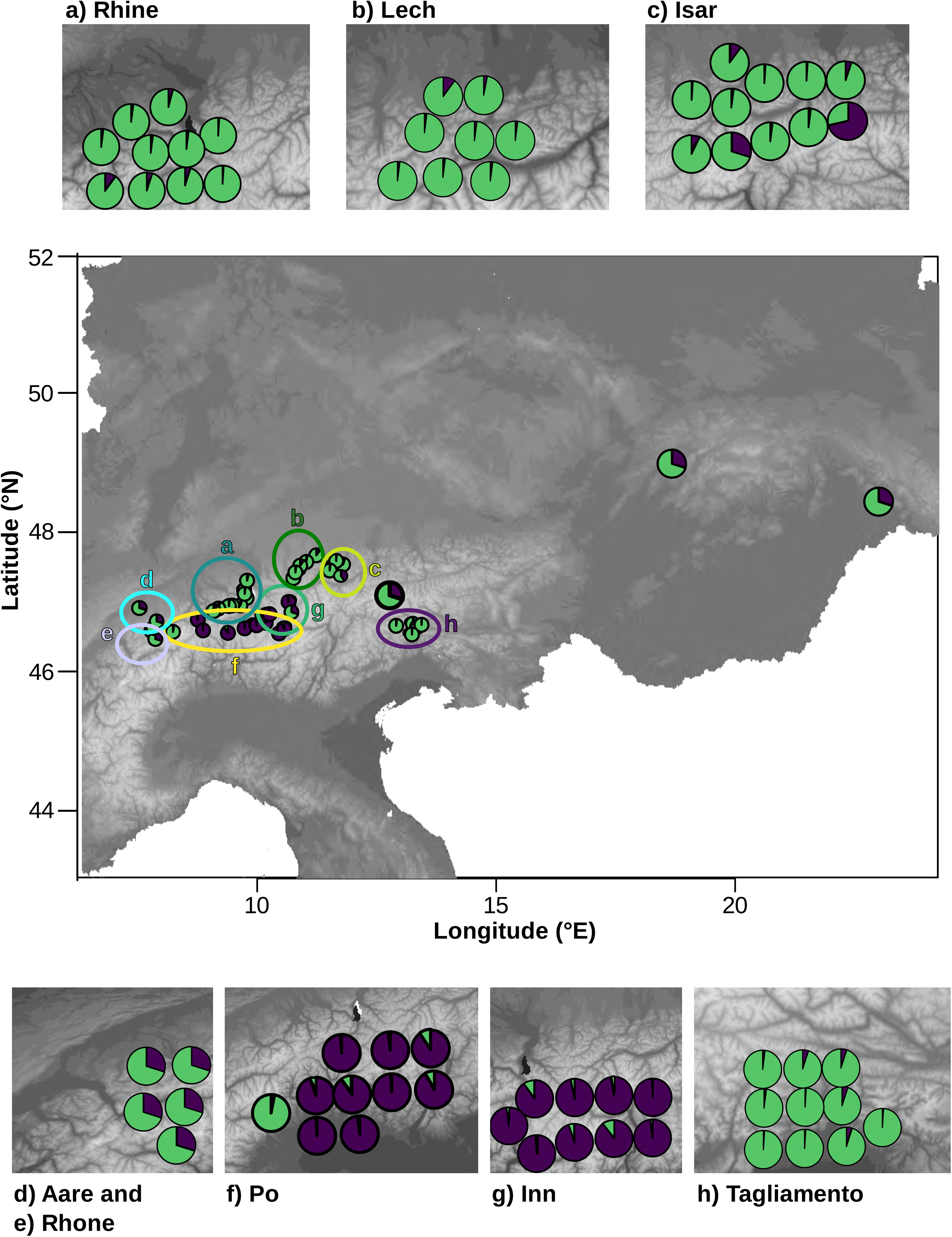
Geographic distribution of nuclear genetic clusters across the 12 river catchments. Pie charts represent the mean assignment proportions (Q values) of the two STRUCTURE clusters (K = 2) for each population. Catchments with a single sampled population are shown with enlarged pie charts for clarity.

STRUCTURE and DAPC analyses were also conducted separately for each of the 12 river catchments (Table 1). In most cases, catchment-level analyses revealed patterns consistent with the global K = 2 structure (Figs. S3–S15).

Pairwise nuclear *F*_ST_ values were moderate to high and increased with geographic distance. Populations separated by large distances (e.g. relative to Cze1 and Ukr1) showed very high differentiation (*F*_ST_ = 0.75–0.95; Fig. S1A). Isolation by distance was tested using a Mantel test comparing pairwise genetic and geographic distances. We detected a significant positive relationship between genetic and geographic distance based on nuclear microsatellites (Mantel’s r = 0.37, p = 0.00001), indicating that geographic distance acts as a barrier to gene flow among populations (Fig. S16). Analysis of molecular variance (AMOVA) attributed 49% of the total variation to differences among populations and 51% to differences among individuals within populations (p < 0.05; Table S3).

#### 3.2.2 Chloroplast microsatellites

STRUCTURE analysis of chloroplast microsatellites also supported two main genetic clusters (K = 2; Fig. 5A–B; Fig. S17), largely concordant with the nuclear results.

**FIGURE 4.**
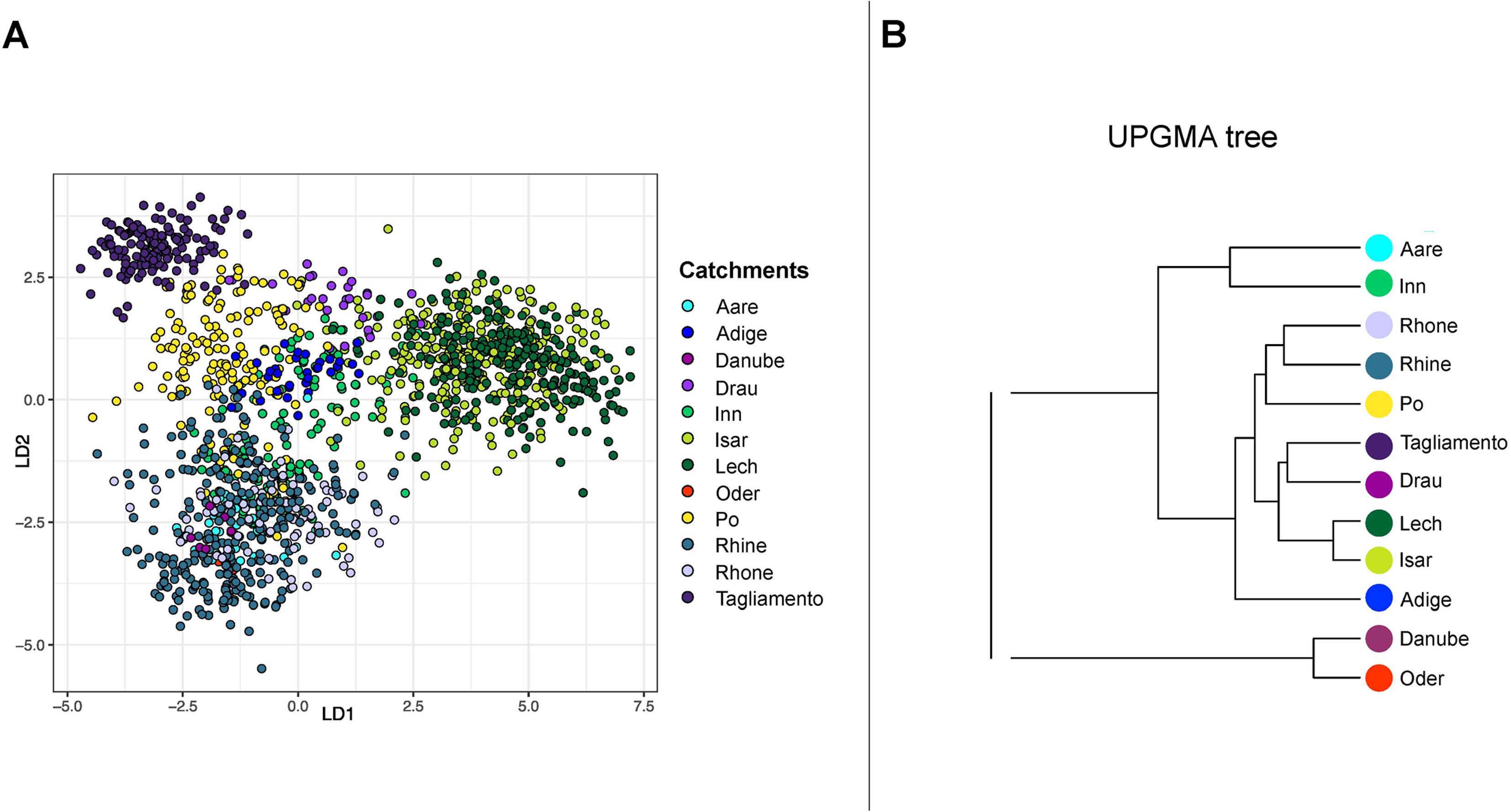
Population genetic structure *M. germanica* inferred from nuclear microsatellites. (A) DAPC scatterplot of individuals coloured by catchment. The first two discriminant axes explain 82% of the total discriminant variance. (B) Unrooted maximum-likelihood tree of catchments based on nuclear genetic distances; bootstrap support (5000 replicates) is shown.

**FIGURE 5.**
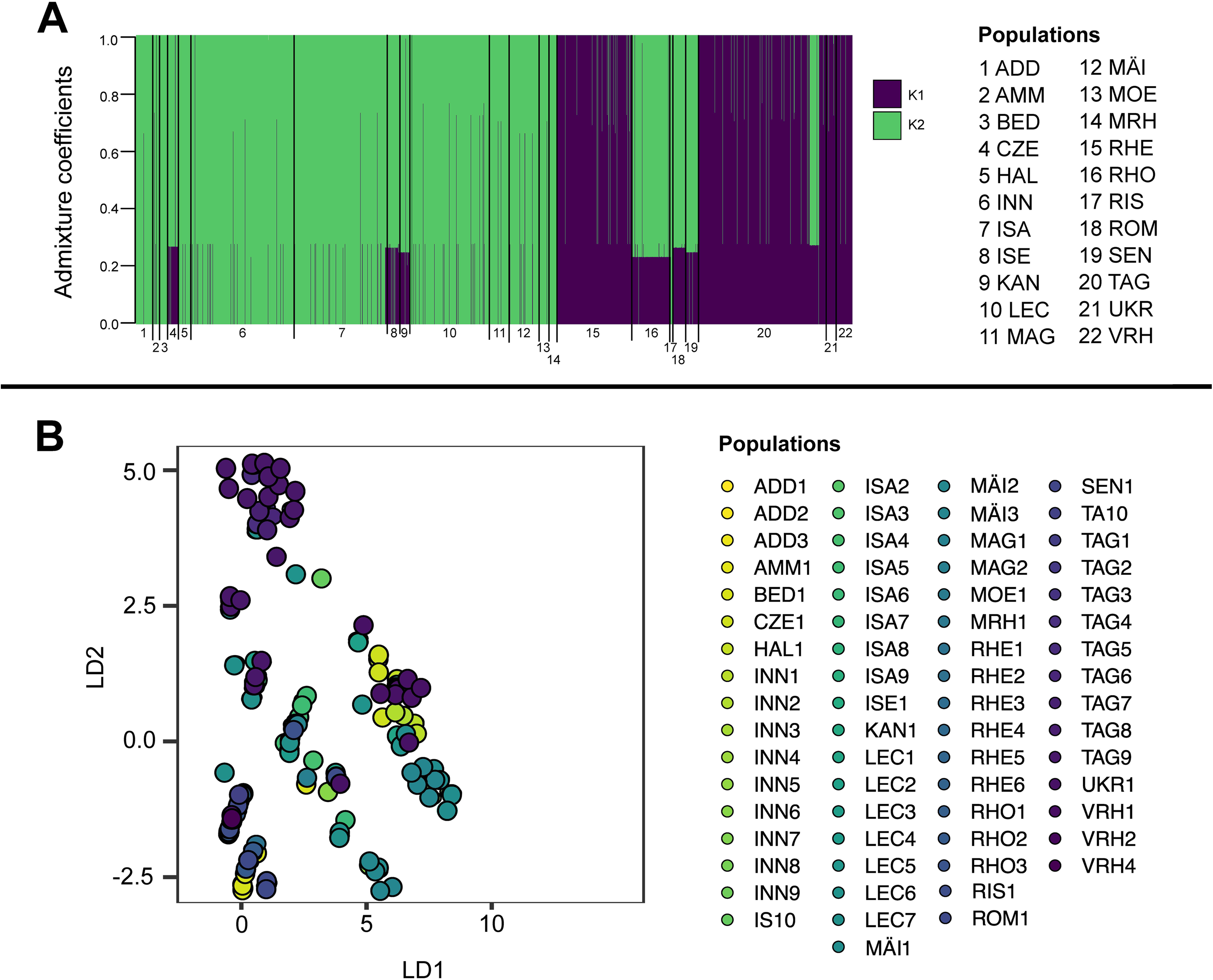
Population genetic structure of *M. germanica* based on six chloroplast microsatellite loci. (A) STRUCTURE clustering performed in haploid mode. Each vertical bar represents an individual, and colours indicate cluster membership coefficients for K = 2. Results are shown descriptively to illustrate chloroplast lineage structure. (B) DAPC scatterplot of individuals coloured by population. The first two discriminant axes explain 90% of the total discriminant variance.

However, chloroplast data showed substantially lower admixture, with most populations dominated by a single cluster (Fig. 5A). Pairwise chloroplast *F*_ST_ values were moderate to high and increased with geographic distance (Fig. S1B), indicating stronger population structure than observed for nuclear markers. DAPC based on chloroplast microsatellites revealed clearer spatial separation among populations compared to nuclear data (Fig. 5B). For chloroplast microsatellites, isolation by distance was weaker but still significant (Mantel’s r = 0.22, p = 0.001), suggesting more complex or historically contingent patterns of seed-mediated dispersal. AMOVA based on chloroplast microsatellite haplotypes revealed strong population structure, with 75.25% of the genetic variation occurring among populations and 24.75% within populations (ΦST ≈ 0.75, p < 0.01), consistent with highly restricted seed-mediated gene flow.

### 3.3 | Genetic structure and dispersal among catchments

#### 3.3.1 Nuclear and chloroplast structure among catchments

DAPC of nuclear microsatellites showed that the first two discriminant axes explained 33.7% of the total genetic variance (Fig. 4A). Individuals grouped into three main clusters: a central cluster containing individuals from most catchments, a second cluster dominated by individuals from the Isar and Lech catchments, and a third cluster comprising individuals from Tagliamento and subsets of Po and Drau.

Clusters two and three were more closely related to each other than to the central group, as reflected by their proximity in the UPGMA tree (Fig. 4B). Populations within the same catchment generally exhibited lower differentiation; however, several cases of high within-catchment *F*_ST_ were observed (Fig. S1A; Figs. S3–S15). Notably, populations exhibiting strong genetic differentiation also showed high inbreeding coefficients, but were not consistently more geographically isolated than other populations within the same catchment (Table 2). Analysis of molecular variance (AMOVA) among catchments (n=12) revealed strong and significant genetic structuring among populations. A total of 36.7% of the genetic variation was attributable to differences among populations (*F*_ST_ = 0.367, 95% CI: 0.332–0.399, *p* = 0.001), while 37.0% of the variation occurred among individuals within populations (*F*_IS_ = 0.584, 95% CI: 0.553–0.620, *p* = 0.001). Only 26.3% of the variation was found within individuals (*F*_IT_ = 0.737). These results indicate pronounced differentiation combined with high levels of inbreeding among catchments.

For chloroplast microsatellites, the first two DAPC axes explained 56.5% of the total variance (Fig. 6A). Individuals again formed three main groups, with stronger separation among catchments than observed for nuclear markers. Nevertheless. some individuals showed mixed ancestry, indicating limited but detectable seed-mediated gene flow. Clusters were also supported by UPGMA analyses (Fig. 6B). AMOVA based on chloroplast microsatellites revealed strong population structure, with the majority of genetic variation occurring among catchments (ΦST = 0.78, p < 0.01), consistent with restricted seed-mediated gene flow. In contrast to nuclear markers, within-population variation was comparatively low, highlighting the stronger spatial structuring of maternally inherited variation.

**FIGURE 6.**
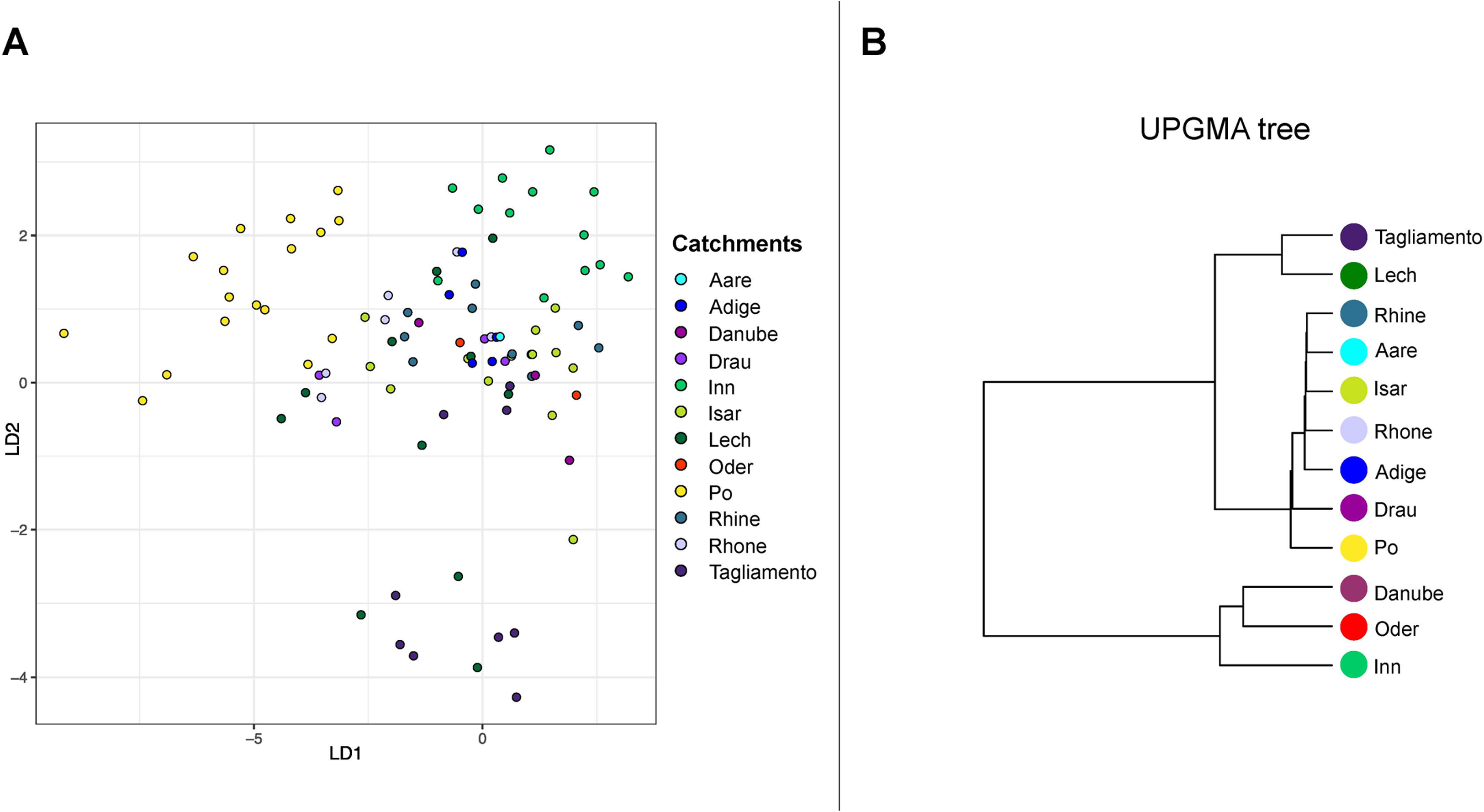
Population genetic structure of *M. germanica* inferred from chloroplast microsatellites. (A) DAPC scatterplot of individuals coloured by catchment. The first two discriminant axes explain 88% of the total discriminant variance. (B) Unrooted maximum-likelihood tree of catchments based on chloroplast genetic distances; bootstrap support (5000 replicates) is shown.

#### 3.3.2 Pollen versus seed dispersal among catchments

Estimated migration rates indicated contrasting dispersal patterns for pollen and seed. Mean pollen migration among catchments was low (mean = 0.06; range = 0.01–0.21), whereas mean seed migration was substantially higher (mean = 0.30; range = 0.14–0.41). The pollen-to-seed migration ratio (mp/ms = 0.136) suggests that seed dispersal contributes more strongly than pollen dispersal to gene flow among catchments in *M. germanica*.

### 3.4 | Migration patterns

#### 3.4.1 Historical nuclear migration

Historical nuclear migration rates inferred using MIGRATE-n were uniformly high. with posterior median M values ranging from approximately ∼60 to >500 (Appendix 1). These estimates indicate pervasive historical nuclear gene flow, with no population approaching isolation. Directionality remained biologically meaningful; however, relative differences among migration pathways were more informative than absolute M values (Appendix 1; Fig. 7A).

**FIGURE 7.**
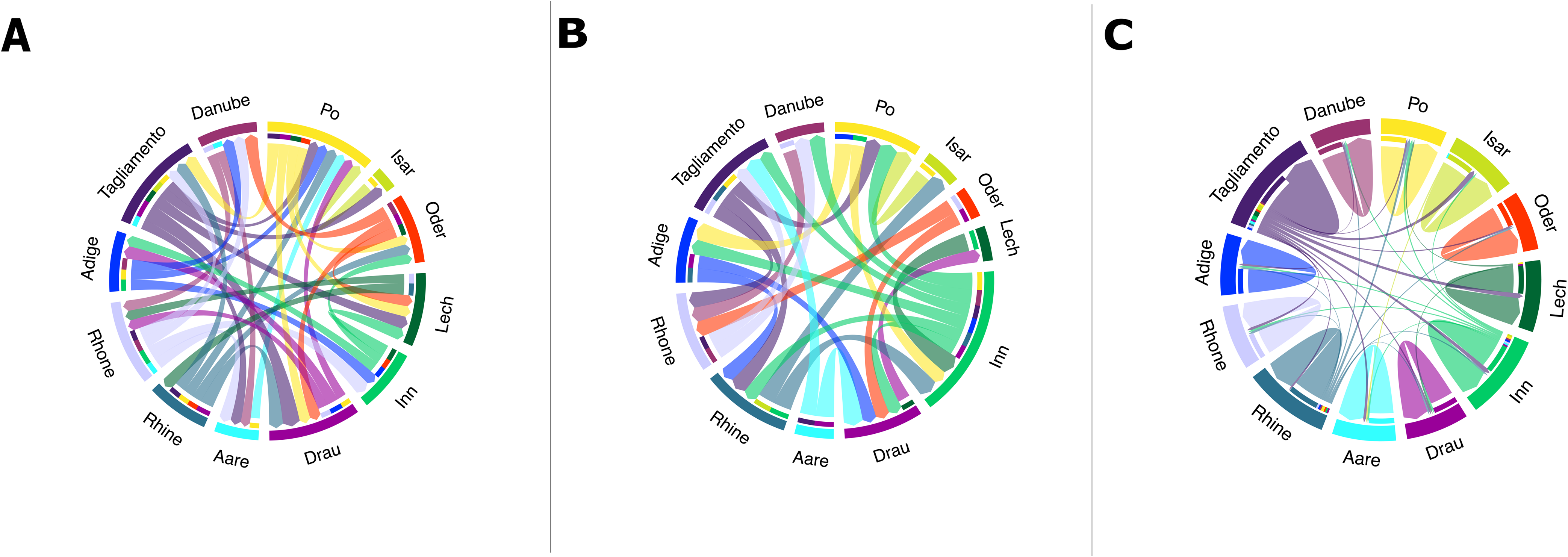
Historical and contemporary migration patterns by seed and pollen dispersal of *M. germanica*. (A) Historical migration among catchments inferred using MIGRATE-N based on nuclear data. Relative migration rates are represented by the proportion of coloured segments within each circle. (B) Historical migration among catchments inferred using MIGRATE-N based on chloroplast haplotypes. Relative migration rates are represented by the proportion of coloured segments within each circle. (C) Contemporary migration among catchments inferred using BayesAss. Migration proportions are represented by coloured segments within each circle.

Several catchments emerged as consistent sinks of nuclear gene flow. The Po catchment received strong immigration from multiple basins, including the Rhine, Aare, Tagliamento, and Adige. Similarly, the Inn catchment showed exceptionally strong incoming gene flow from the Rhone and Adige, indicating pronounced east–west connectivity across Alpine catchments. In contrast, the Rhine and Danube acted as major exporters of nuclear gene flow, with the Danube forming a continental-scale connectivity backbone linking eastern, northern, and southern regions (Appendix 1; Fig. 7A, Table S5).

#### 3.4.2 Historical chloroplast migration

Bayesian estimates of mutation-scaled migration rates based on chloroplast markers revealed generally low but highly directional historical gene flow. Only a subset of migration pathways showed strong posterior support for non-zero migration, while many credible intervals overlapped zero. The strongest supported pathway was migration from the Rhone into the Oder catchment (median M ≈ 52.3; 95% CI: 23.3–78.7; Appendix 2; Fig. 7B, Table S6), followed by migration from the Rhine into Oder (median M ≈ 41.0) and from Lech into Drau (median M ≈ 51.0). Several additional pathways showed moderate support, including migration from Po and Isar into Oder (median M ≈ 30–37)(Appendix 2, Fig. 7B, Table S6).

#### 3.4.3 Contemporary seed migration

Contemporary migration estimated with BA3 revealed low seed-mediated migration among catchments (mean m ± SD = 0.025 ± 0.006) and genetic exchange occurring almost exclusively within catchments (mean m ± SD = 0.75 ± 0.02). Nonetheless, limited gene flow among clusters was detected (Fig. 7C, Appendix S3, Table S7).

Some catchments contributed disproportionately to the gene pool of *M. germanica*, particularly Tagliamento, Rhine, Lech, Inn, and Isar. Inbreeding coefficients were high across all 12 catchments (mean ± SD = 0.72 ± 0.035; Appendix S3).

## 4 | Discussion

Using nuclear and chloroplast microsatellite markers, we investigated genetic diversity, population structure, and gene flow in *M. germanica* across 67 populations spanning 12 major catchments in Central Europe. Across marker types and analytical approaches, our results consistently indicate low genetic diversity, high inbreeding, strong population differentiation, and restricted contemporary connectivity among catchments. At the same time, historical migration analyses reveal pronounced contrasts between pollen- and seed-mediated dispersal, highlighting how dispersal vectors operating over different temporal scales have jointly shaped the current genetic structure of this riparian pioneer species.

### Genetic diversity, inbreeding, and population structure in a riparian context

The low levels of genetic diversity and high inbreeding coefficients observed across populations are consistent with expectations for riparian pioneer plants that combine self-pollination with frequent demographic turnover (Reed & Frankham 2003; Werth & Scheidegger 2014; Fink et al. 2022). Such species often experience repeated founder events during colonization of newly available habitats, followed by population bottlenecks driven by river dynamics and habitat instability. These processes reduce effective population size and increase inbreeding, even when populations are not strongly geographically isolated.

High genetic differentiation among populations, reflected in elevated pairwise *F*_ST_ values for both nuclear and chloroplast markers, further supports this interpretation. Although isolation by distance was detected at the regional scale, we also observed strong differentiation among populations within the same catchment. This pattern has been reported previously for *M. germanica* and other riparian taxa and likely reflects fine-scale habitat fragmentation imposed by river morphology (e.g. canyons. braided sections) and anthropogenic barriers that disrupt dispersal along river corridors (Werth et al. 2014; Fink et al. 2022). Consequently, genetic structure in *M. germanica* appears to be shaped not only by geographic distance but also by local riverscape configuration.

### Contrasting nuclear and chloroplast structure; dispersal vectors matter

Differences between nuclear and chloroplast genetic structure provide further insight into the mechanisms underlying connectivity. Chloroplast microsatellites revealed stronger population structure and lower admixture than nuclear markers, a pattern widely documented in plant species and typically attributed to the smaller effective population size and uniparental inheritance of organellar genomes (Petit et al. 2005; Zhang & Hewitt 2003). In *M. germanica*, this contrast indicates that seed dispersal is more spatially restricted than pollen dispersal, leading to stronger spatial genetic structure in chloroplast DNA.

At the catchment scale, nuclear data showed clearer signatures of isolation by distance than chloroplast data, whereas chloroplast structure sometimes deviated from simple geographic patterns. This suggests that seed-mediated dispersal is influenced by a combination of hydrological connectivity, historical contingencies, and stochastic colonization events, rather than by geographic distance alone (Nilsson et al. 2010; Werth et al. 2014; Auffret et al. 2015). Such decoupling between pollen- and seed-mediated dispersal has been reported for other riparian and woody plant species (Ennos 1994; Petit et al. 2005; Ngeve et al. 2017) and underscores the importance of explicitly comparing marker systems when interpreting population connectivity.

### Historical versus contemporary gene flow

Migration analyses further emphasize the temporal dimension of gene flow in *M. germanica*. Historical nuclear migration rates inferred with MIGRATE-n were uniformly high, indicating widespread pollen-mediated connectivity across catchments over evolutionary timescales (Beerli & Felsenstein 2001; Ennos 1994). No population showed evidence of long-term isolation, and several catchments consistently acted as major sources or sinks of gene flow. In particular, the Rhine and Danube emerged as strong exporters of nuclear gene flow, while the Po and Inn acted as prominent recipients, pointing to structured but highly connected historical networks rather than panmixia (Tockner & Stanford 2002; Burkhardt-Holm et al. 2002; Weiss et al. 2018).

In contrast, chloroplast migration estimates revealed fewer, more directional historical seed dispersal routes, with strong support for only a limited number of pathways. This indicates that seed-mediated gene flow has historically occurred via rare but consequential dispersal events, likely associated with major hydrological connections (Hewitt 2000. 2004; Nilsson et al. 2010). Contemporary migration analyses based on BA3 reinforce this view, showing that current seed-mediated gene flow among catchments is very low, with genetic exchange occurring almost exclusively within catchments (Wilson & Rannala 2003; Werth & Scheidegger 2014). Thus, while historical pollen flow has maintained nuclear cohesion across large spatial scales, present-day seed dispersal is strongly constrained by catchment boundaries.

### Alpine glaciation and postglacial recolonization

The contrasting spatial signatures of nuclear and chloroplast gene flow are consistent with the legacy of Alpine glaciation and postglacial recolonization. During Pleistocene glacial maxima, *M. germanica* was likely restricted to low-elevation refugia associated with major river valleys and foreland basins (Taberlet et al. 1998; Hewitt 2000; Schönswetter et al. 2005). Postglacial expansion from these refugia would have proceeded primarily along river corridors, generating strong directional signals in maternally inherited chloroplast markers (Hewitt 2004; Nilsson et al. 2010).

The pronounced chloroplast structure and the concentration of seed-mediated gene flow into specific catchments suggest recolonization via a limited number of dispersal routes rather than widespread, multidirectional expansion. In contrast, the extensive historical nuclear connectivity likely reflects prolonged pollen-mediated gene flow following recolonization, which progressively eroded early colonization signals in the nuclear genome while leaving chloroplast structure largely intact (Ennos 1994; Petit et al. 2005). The identification of the Rhine and Danube as major exporters of nuclear gene flow is consistent with their role as long-term demographic reservoirs and postglacial dispersal corridors facilitating secondary contact among lineages (Tockner & Stanford 2002; Burkhardt-Holm et al. 2002; Weiss et al. 2018).

Together, these results support a two-phase postglacial history in *M. germanica*; an initial phase dominated by rare, directional seed dispersal shaping chloroplast structure, followed by extended pollen-mediated gene flow that maintained nuclear connectivity across catchments.

### Conservation implications

From a conservation perspective, our findings highlight catchments as biologically meaningful management units for *M. germanica*, particularly with respect to seed-mediated dispersal and local demographic processes. Contemporary gene flow is largely confined within catchments, and high inbreeding coefficients across populations suggest limited effective dispersal and strong local mating. Disruption of within-catchment connectivity through river regulation, habitat fragmentation, or land-use change is therefore likely to have immediate negative consequences for population viability (Fausch et al. 2002; Bunn & Arthington 2002).

At the same time, the persistence of historical pollen-mediated connectivity underscores the importance of maintaining landscape permeability at broader spatial scales. Large catchments such as the Rhine and Danube appear to function as genetic reservoirs and sources of connectivity, while catchments such as the Po and Inn accumulate genetic diversity through admixture. Conservation strategies should thus prioritize both the preservation of within-catchment habitat continuity and the maintenance of regional connectivity networks that facilitate long-term genetic exchange.

## 5 | Conclusions

In summary, *M. germanica* populations across Central Europe are characterized by low genetic diversity, high inbreeding, and strong spatial structure, shaped by limited seed dispersal, selfing, and dynamic riverscape processes. Despite this fine-scale structure, historical pollen-mediated gene flow has generated widespread nuclear connectivity among catchments, reflecting the enduring influence of postglacial dispersal corridors. The pronounced mismatch between historical and contemporary connectivity highlights the vulnerability of riparian species to ongoing hydrological alteration and habitat fragmentation. By integrating multiple marker systems and temporal perspectives, our study demonstrates that *M. germanica* is best described as structured but connected, and emphasizes the need for catchment-based conservation strategies that preserve both local demographic processes and long-term evolutionary connectivity.

## ACKNOWLEDGMENTS

Logistic support was received from Genetic Diversity Centre (GDC) of ETH Zürich. From WSL. From University of Iceland and from LMU Munich. We thank A. Minder and T. Torrossi (GDC) for running fragment analyses on an automatic sequencer. The data generation was kindly funded by the Swiss Federal Office of the Environment. BAFU for the research program Hydraulic Engineering and Ecology. Barbara Krummenacher collected many of the samples included in this study. We thank S. Angelone. T. Karpati and C. Spinelli for helping with the fieldwork. S. Cheenacharoen and Y. Kophimai helped with extracting DNA.

## AUTHOR CONTRIBUTIONS

Conceived and designed the experiments: SW. CS. Performed the experiments: SW. Generated all data: SW. Analyzed the data: SW. TCP. Made the figures: TCP.

Improved the figures: KS. Wrote the paper: TCP. Edited the manuscript: CS. KS. SW. TCP.

## OPEN RESEARCH BADGES

### DATA AVAILABILITY STATEMENT

The nuclear loci and the chloroplast loci data. The population and individual information of the samples and the R script have been deposited under the following link: https://figshare.com/s/13083619cea77f51ab22

