## Supplementary material for "River network connectivity and postglacial history shape pollen- and seed-mediated gene flow across riparian populations of *Myricaria germanica*": Supplementary Information Myricaria paper Chavarria-Pizarro et al 2026..docx

This file includes:

Figs. S1- S18

Tables S1-S2

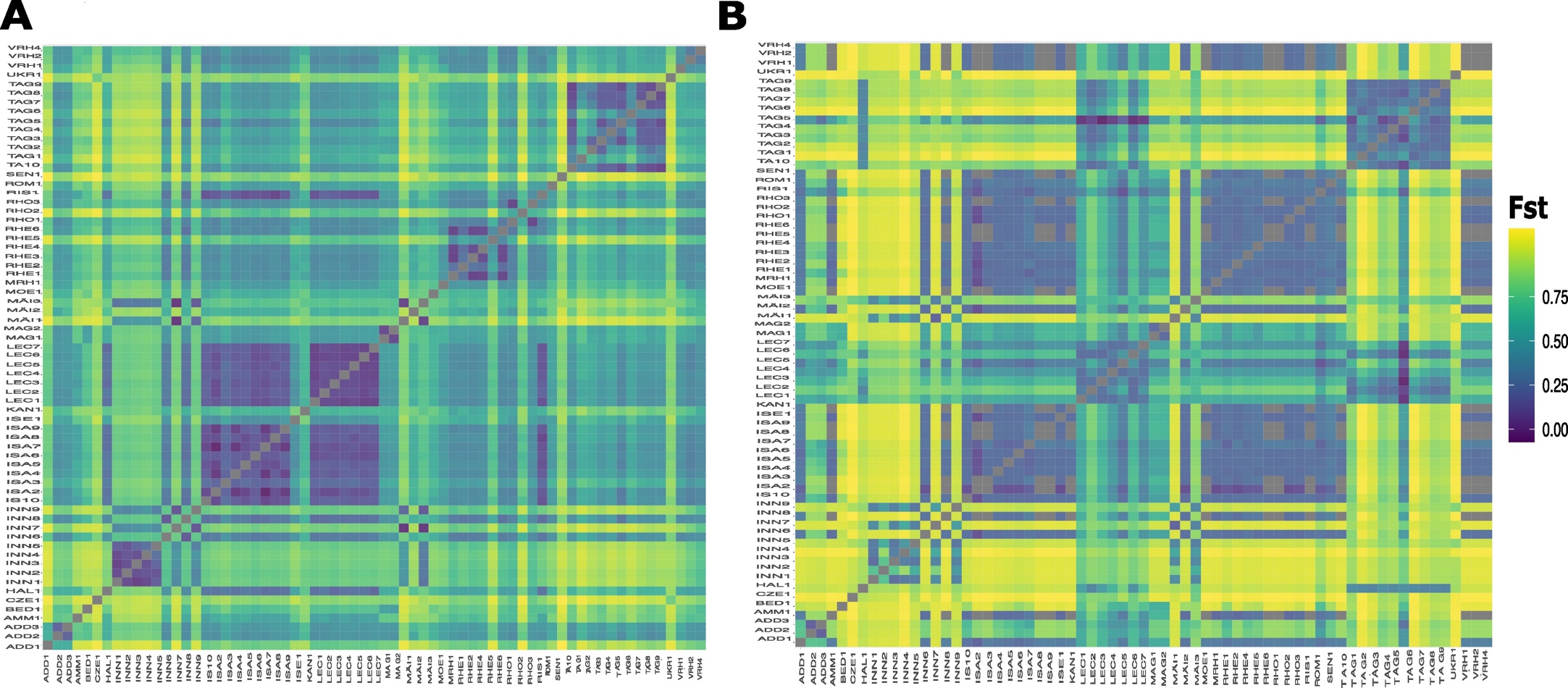

S1. Heatmap with population pairwise *F*_ST_ values (0-1) among 67 populations of *M. germanica*. Lower values are dark purple and higher values are yellow. A) Heatmap of population pairwise *F*_ST_ values (0-1) based on 20 nuclear microsatellite data B) Heatmap of population pairwise *F*_ST_ values (0-1) based on 6 chloroplast microsatellite data.

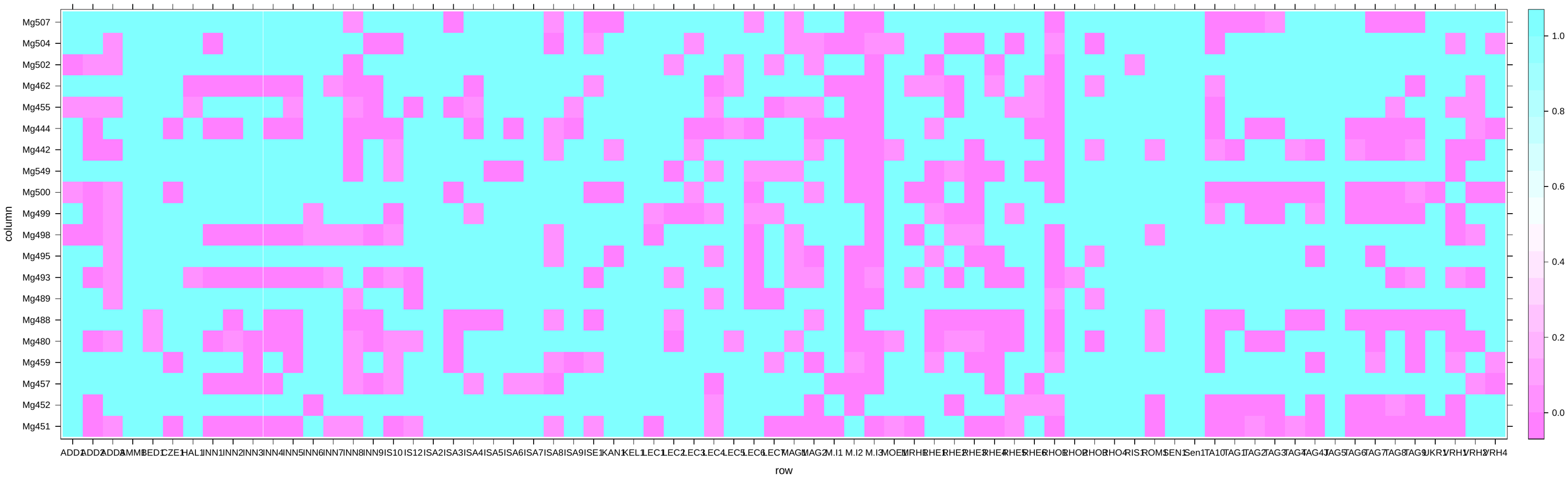

S2. Heatmap of Hardy-Weinberg disequilibrium values base on 20 nuclear micrpsatellite data, ranging from 0 to 1, the colors represent the p-values, where pinker colors indicate values closer to 0 (significant disequilibrium) and bluer colors indicate values closer to 1 (in equilibrium).

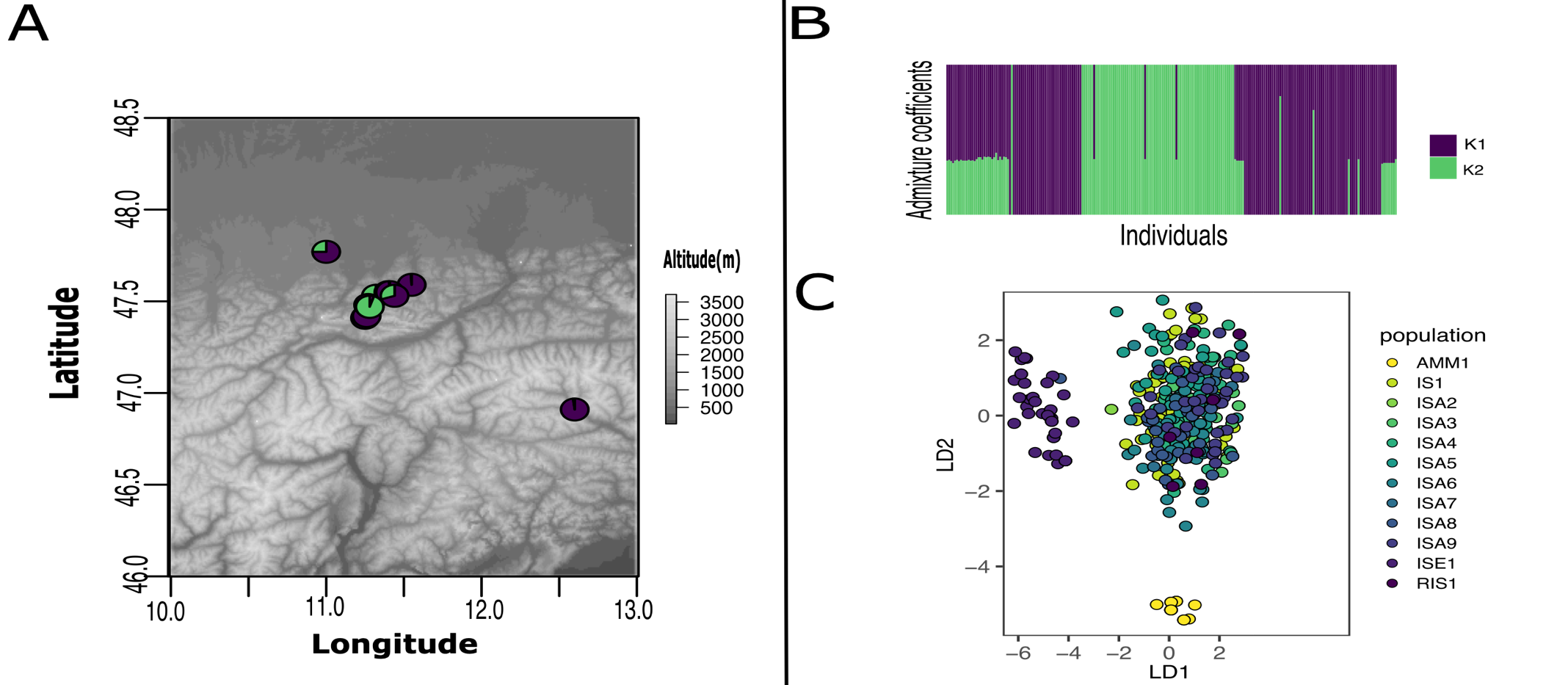

S3. Population genetic structure of *Myricaria germanica* of Isar catchment populations based on 20 nuclear microsatellites A) Geographic distribution and lineage assignments of 12 populations of Isar catchment, K1 is dark purple and K2 is dark green. The diagram represents the average of each proportion of assignment (Q) of the genetic groups for the population contemporary gene pool. B) Graph of cluster analysis in STRUCTURE. Each horizontal bar represents an individual. The colors represent the coefficient of association for each genetic group base on the ΔK statistic likelihood function which identified K = 2 as the most appropriate number of genetic groups. K1 is dark purple and K2 is dark green C) Scatterplot of the genetic structure of discriminant analysis of principal components (DAPC) showing the individuals (points) from color-coding of the 12 populations, the first two axes that together explained 83% of the total variance, and individuals from all areas were somehow separated in few small groups.

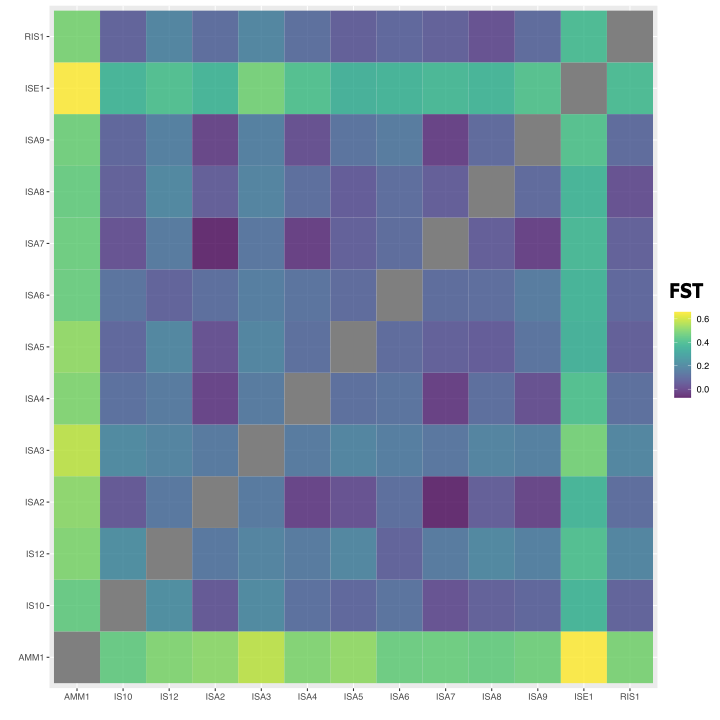

S4. Heatmap with population pairwise *F*_ST_ values (0-1) based on 20 nuclear microsatellite data among 12 populations of *M. germanica* on Isar catchment. Lower values are dark purple and higher values are yellow.

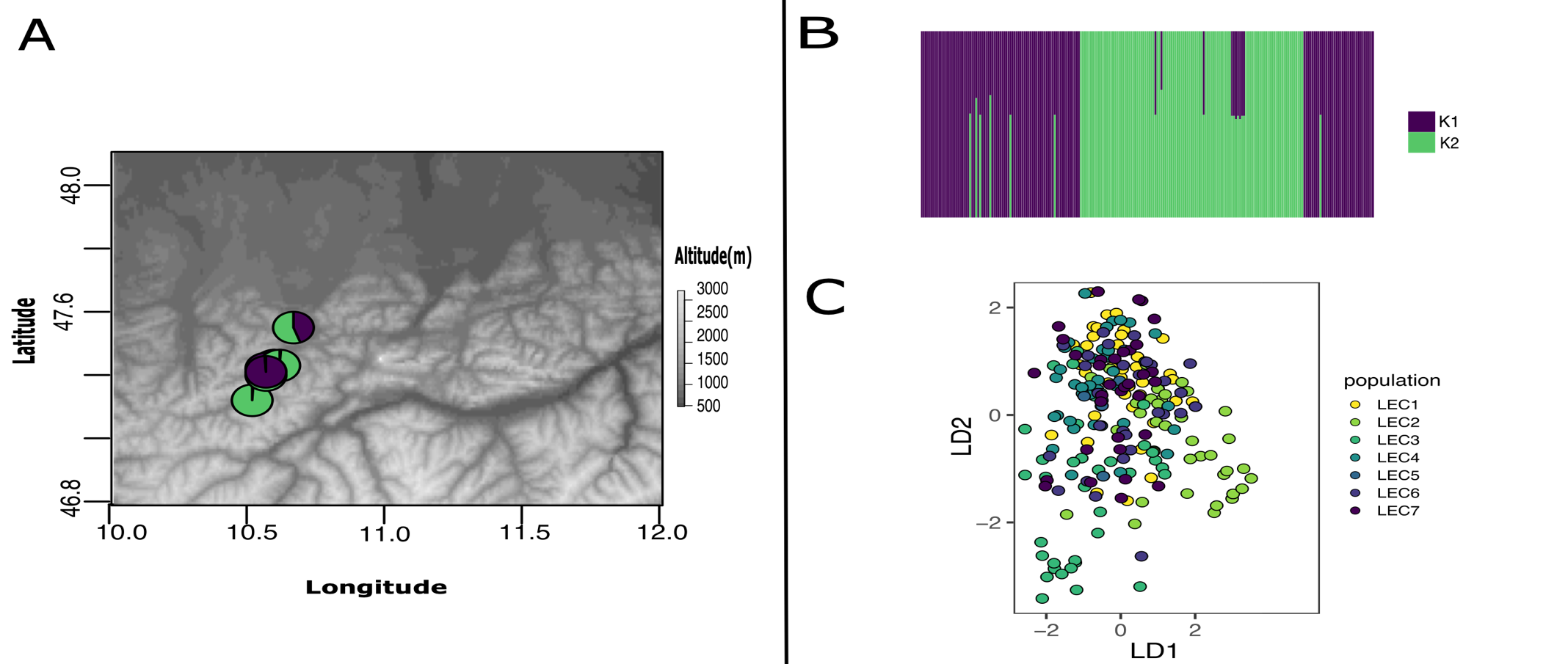

S5. Population genetic structure of *Myricaria germanica* of Lech catchment populations based on 20 nuclear microsatellites A) Geographic distribution and lineage assignments of 7 populations of Lech catchments, K1 is dark purple and K2 is dark green. The diagram represents the average of each proportion of assignment (Q) of the genetic groups for the population contemporary gene pool. B) Graph of cluster analysis in STRUCTURE. Each horizontal bar represents an individual. The colors represent the coefficient of association for each genetic group base on the ΔK statistic likelihood function which identified K = 2 as the most appropriate number of genetic groups. K1 is dark purple and K2 is dark green C) Scatterplot of the genetic structure of discriminant analysis of principal components (DAPC) showing the individuals (points) from color-coding of the 7 populations, the first two axes that together explained 65% of the total variance, and individuals from all areas were somehow separated in few small groups.

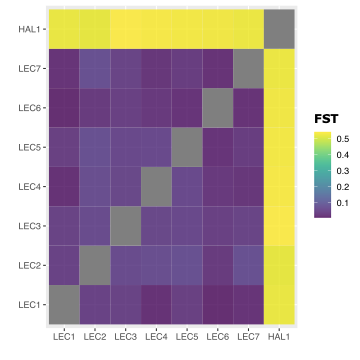

S6. Heatmap with population pairwise *F*_ST_ values (0-1) based on 20 nuclear microsatellite data among 7 populations of *M. germanica* on Lech catchment. Lower values are dark purple and higher values are yellow.

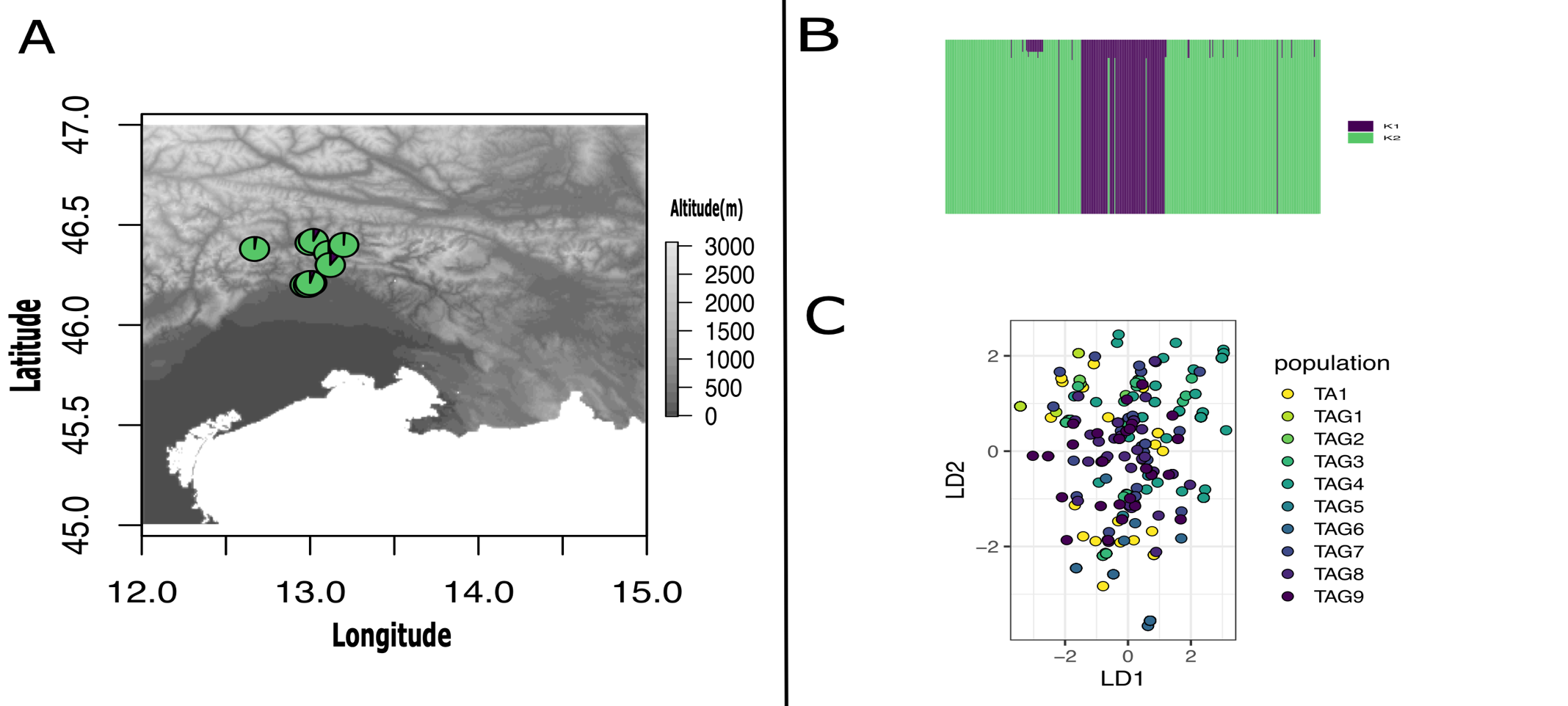

S7. Population genetic structure of *M. germanica* of Taglimento catchment populations, based on 20 nuclear microsatellites A) Geographic distribution and lineage assignments of 10 populations of Taglimento catchment, K1 is dark purple and K2 is dark green. The diagram represents the average of each proportion of assignment (Q) of the genetic groups for the population contemporary gene pool. B) Graph of cluster analysis in STRUCTURE. Each horizontal bar represents an individual. The colors represent the coefficient of association for each genetic group base on the ΔK statistic likelihood function which identified K = 2 as the most appropriate number of genetic groups. K1 is dark purple and K2 is dark green C) Scatterplot of the genetic structure of discriminant analysis of principal components (DAPC) showing the individuals (points) from color-coding of the 10 populations, the first two axes that together explained 73% of the total variance, and individuals from all areas were somehow separated in few small groups.

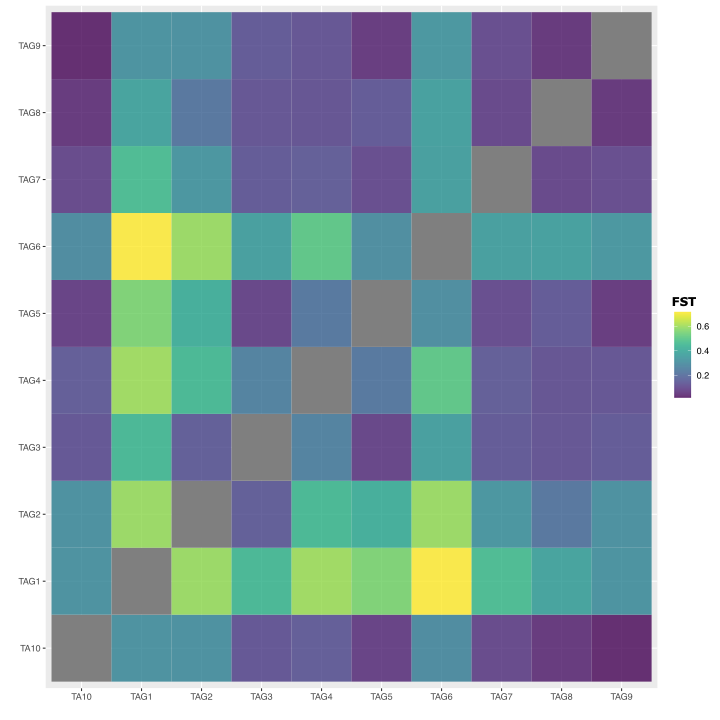

S8. Heatmap with population pairwise *F*_ST_ values (0-1) based on 20 nuclear microsatellite data among 7 populations of *M. germanica* on Taglimento catchment. Lower values are dark purple and higher values are yellow.

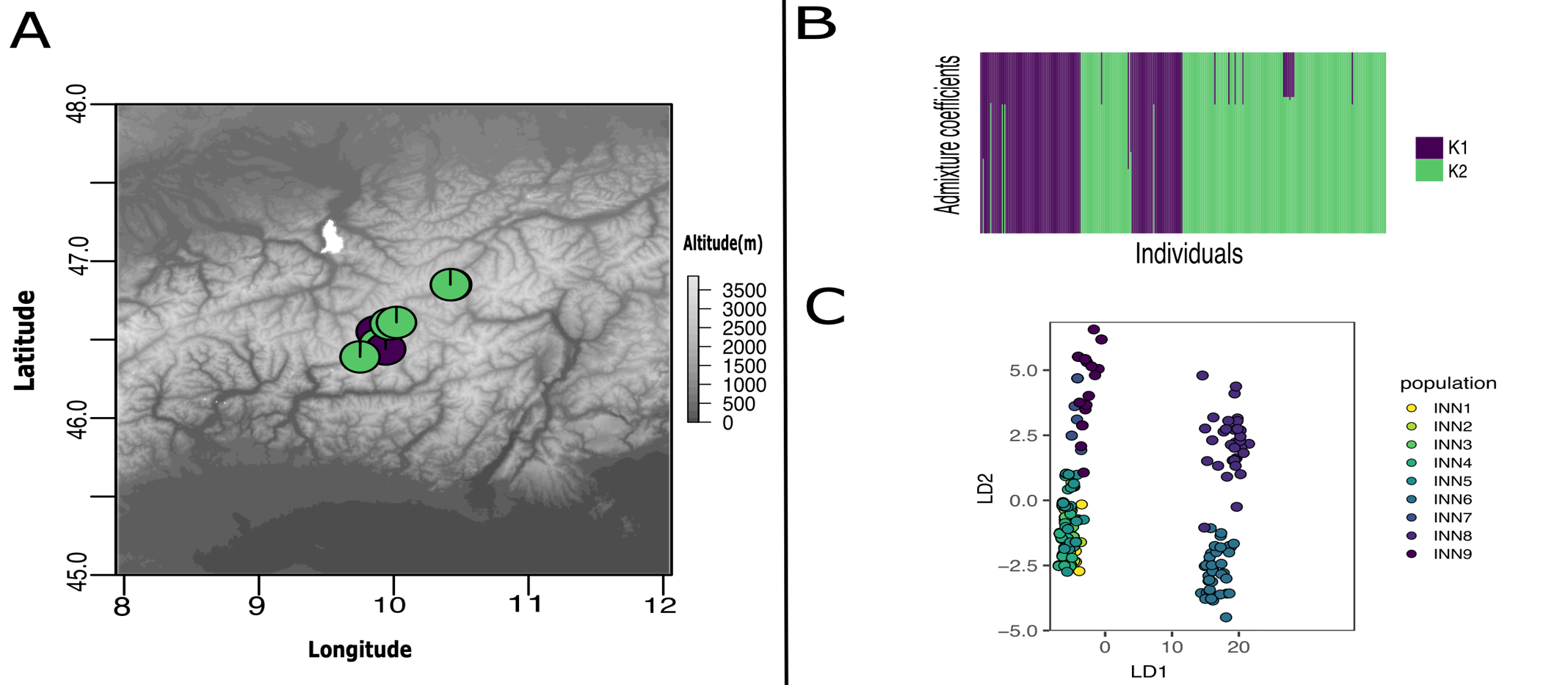

S9. Population genetic structure of *M. germanica* of Inn catchment populations, based on 20 nuclear microsatellites A) Geographic distribution and lineage assignments of 9 populations of Inn catchment, K1 is dark purple and K2 is dark green. The diagram represents the average of each proportion of assignment (Q) of the genetic groups for the population contemporary gene pool. B) Graph of cluster analysis in STRUCTURE. Each horizontal bar represents an individual. The colors represent the coefficient of association for each genetic group base on the ΔK statistic likelihood function which identified K = 2 as the most appropriate number of genetic groups. K1 is dark purple and K2 is dark green C) Scatterplot of the genetic structure of discriminant analysis of principal components (DAPC) showing the individuals (points) from color-coding of the 9 populations, the first two axes that together explained 88% of the total variance, and individuals from all areas were somehow separated in few small groups.

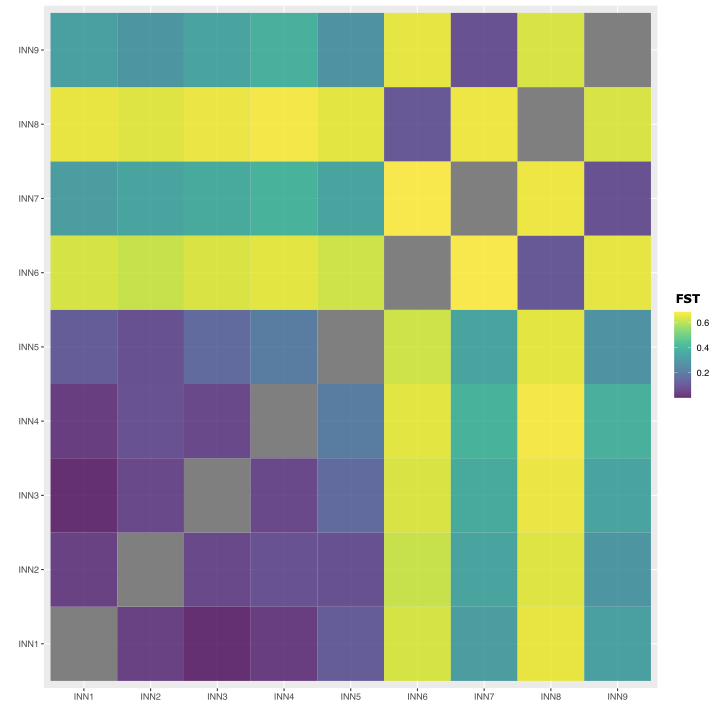

S10. Heatmap with population pairwise *F*_ST_ values (0-1) based on 20 nuclear microsatellite data among 9 populations of *M. germanica* on Inn catchment. Lower values are dark purple and higher values are yellow.

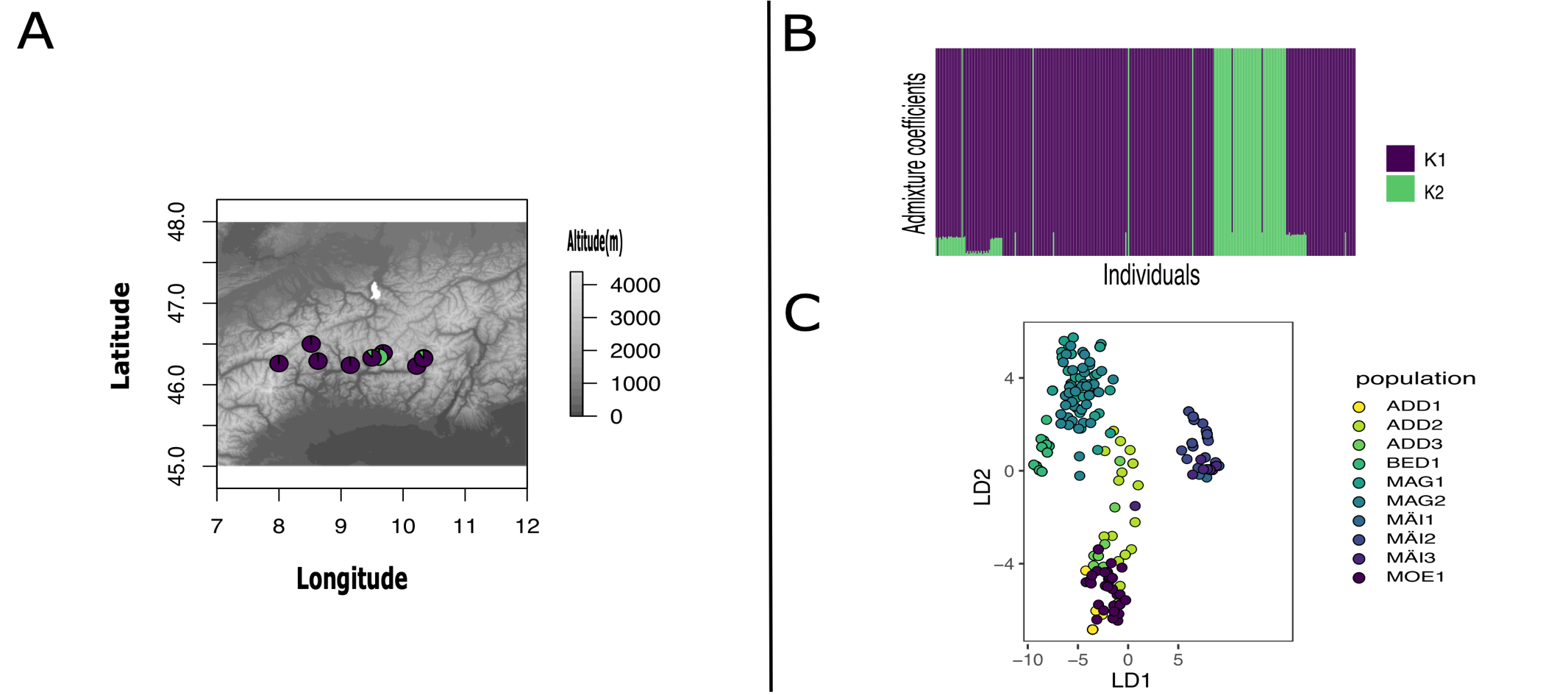

S11. Population genetic structure of *M. germanica* of Po catchment populations, based on 20 nuclear microsatellites A) Geographic distribution and lineage assignments of 10 populations of Po catchment, K1 is dark purple and K2 is dark green. The diagram represents the average of each proportion of assignment (Q) of the genetic groups for the population contemporary gene pool. B) Graph of cluster analysis in STRUCTURE. Each horizontal bar represents an individual. The colors represent the coefficient of association for each genetic group base on the ΔK statistic likelihood function which identified K = 2 as the most appropriate number of genetic groups. K1 is dark purple and K2 is dark green C) Scatterplot of the genetic structure of discriminant analysis of principal components (DAPC) showing the individuals (points) from color-coding of the 10 populations, the first two axes that together explained 91% of the total variance, and individuals from all areas were somehow separated in few small groups.

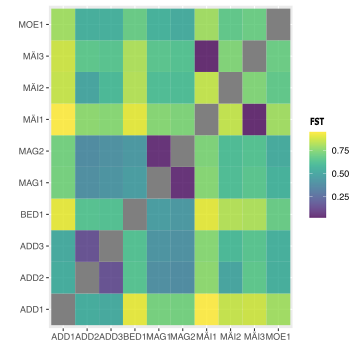

S12. Heatmap with population pairwise *F*_ST_ values (0-1) based on 20 nuclear microsatellite data among 10 populations of *M. germanica* on Po catchment. Lower values are dark purple and higher values are yellow.

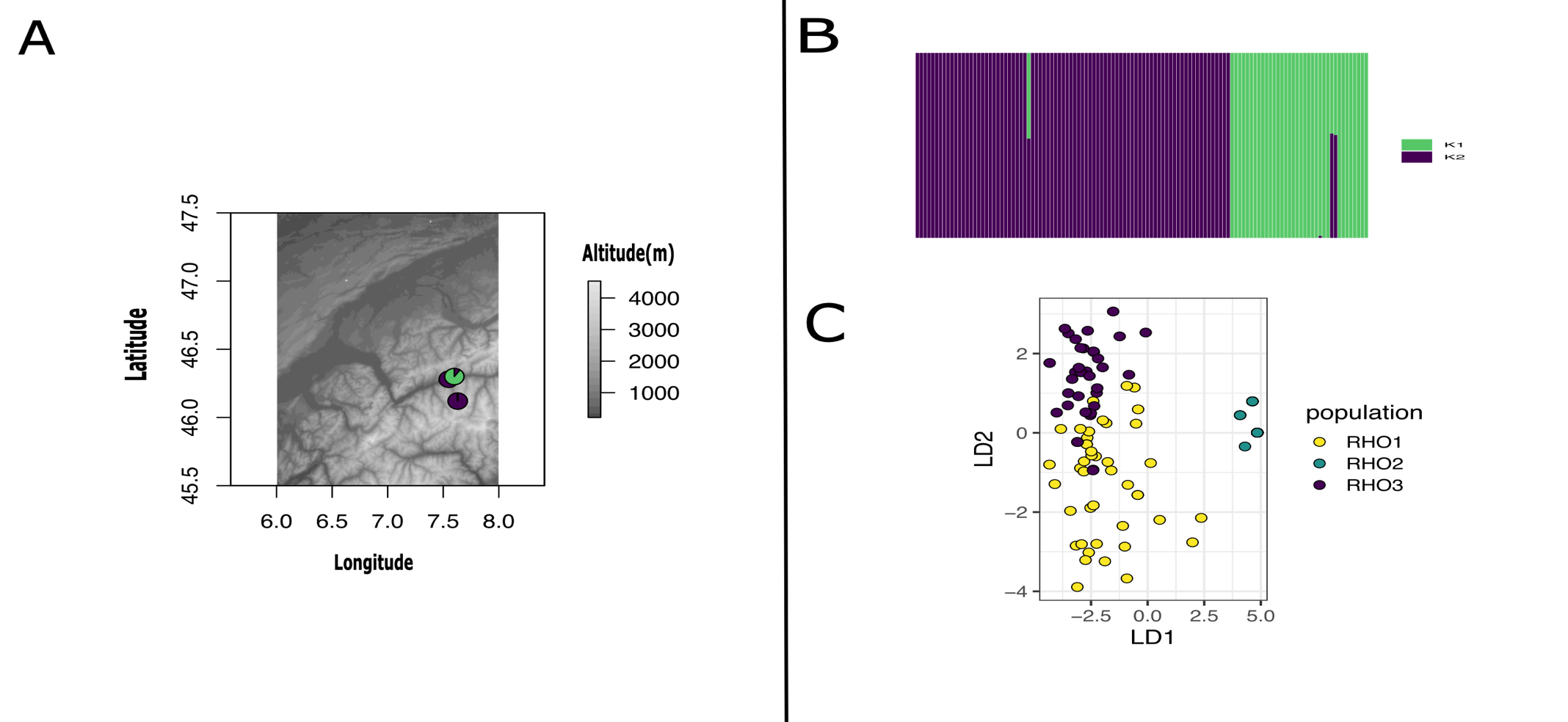

S13. Population genetic structure of *M. germanica* of Rhone catchment populations, based on 20 nuclear microsatellites A) Geographic distribution and lineage assignments of 3 populations of Rhone catchment, K1 is dark purple and K2 is dark green. The diagram represents the average of each proportion of assignment (Q) of the genetic groups for the population contemporary gene pool. B) Graph of cluster analysis in STRUCTURE. Each horizontal bar represents an individual. The colors represent the coefficient of association for each genetic group base on the ΔK statistic likelihood function which identified K = 2 as the most appropriate number of genetic groups. K1 is dark purple and K2 is dark green C) Scatterplot of the genetic structure of discriminant analysis of principal components (DAPC) showing the individuals (points) from color-coding of the 3 populations, the first two axes that together explained 95% of the total variance, and individuals from all areas were somehow separated in few small groups.

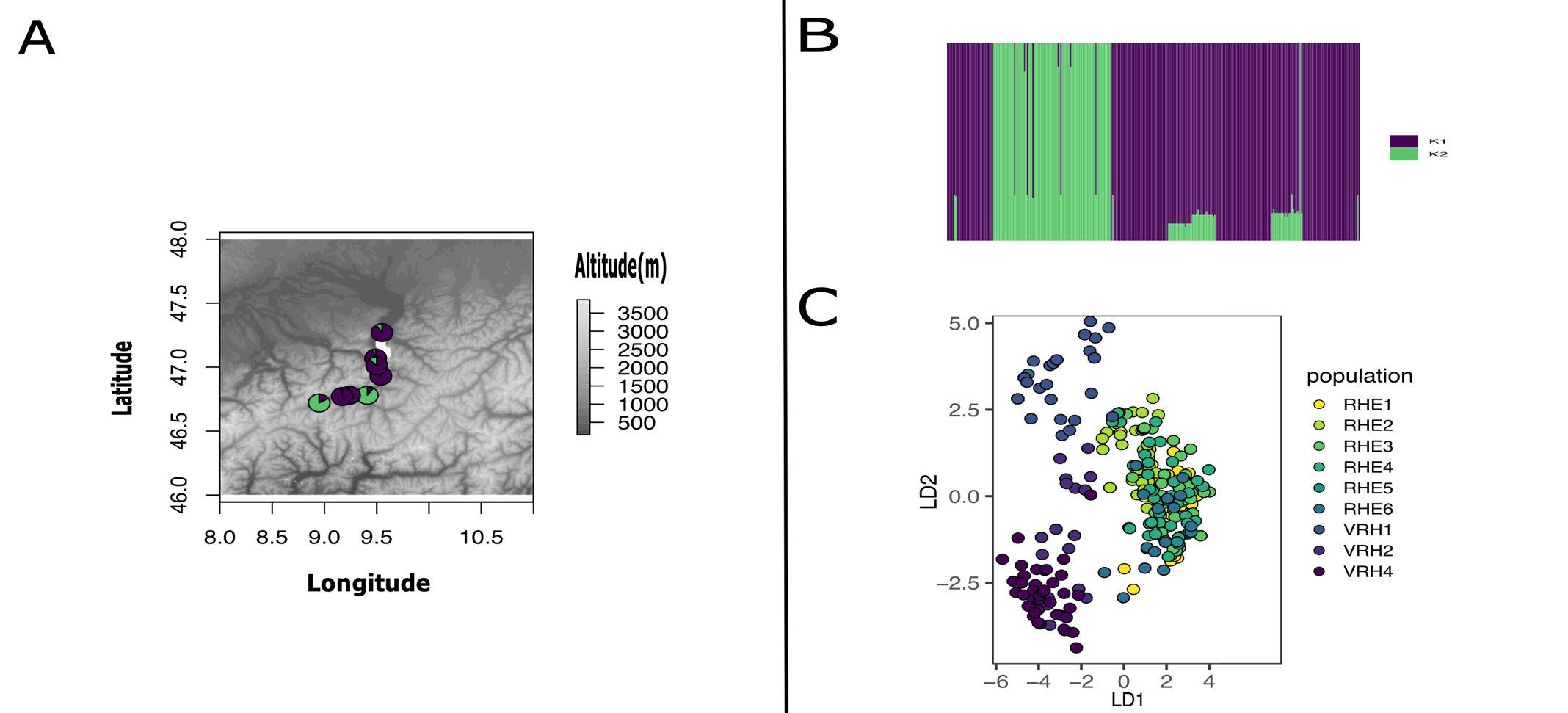

S14. Population genetic structure of *M. germanica* of Rhine catchment populations, based on 20 nuclear microsatellites A) Geographic distribution and lineage assignments of 9 populations of Rhine catchment, K1 is dark purple and K2 is dark green. The diagram represents the average of each proportion of assignment (Q) of the genetic groups for the population contemporary gene pool. B) Graph of cluster analysis in STRUCTURE. Each horizontal bar represents an individual. The colors represent the coefficient of association for each genetic group base on the ΔK statistic likelihood function which identified K = 2 as the most appropriate number of genetic groups. K1 is dark purple and K2 is dark green C) Scatterplot of the genetic structure of discriminant analysis of principal components (DAPC) showing the individuals (points) from color-coding of the 9 populations, the first two axes that together explained 89% of the total variance, and individuals from all areas were somehow separated in few small groups.

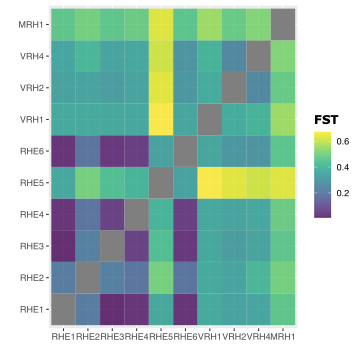

S15. Heatmap with population pairwise *F*_ST_ values (0-1) based on 20 nuclear microsatellite data among 9 populations of *M. germanica* on Rhine catchment. Lower values are dark purple and higher values are yellow.

S16. Scatterplot showing geographic (km) vs genetic distance (*F*_ST_) among 67 populations of *M.germanica* based on 20 nuclear loci. Mantel test revealed a positive significant correlation between geographic and genetic distance (Mantel’s *r* = .37, *p* = .00001), demonstrating that geographical distance represents a barrier to gene flow for *M. germanica.*

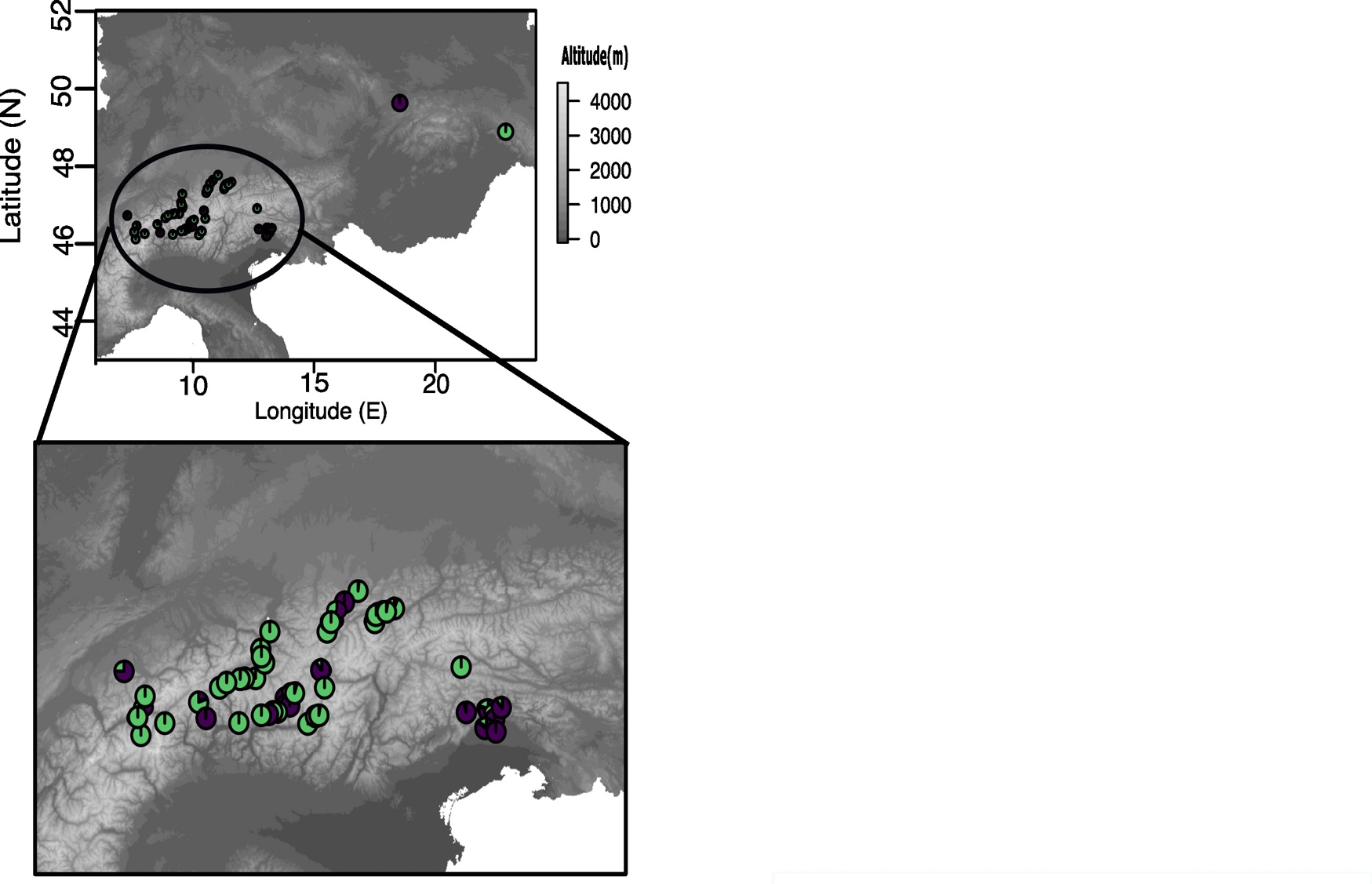

S17. Geographic distribution and lineage assignments of *M. germanica* based on 6 chloroplast loci of the 67 populations from the 12 catchments. K1 is dark purple and K2 is dark green. The diagram represents the average of each proportion of assignment (Q) of the genetic groups for the population historical gene pool.

S18. Scatterplot showing geographic (km) vs genetic distance (*F*_ST_) among 67 populations of *M.germanic*a based on 6 chloroplast loci. Mantel test revealed a positive significant correlation between geographic and genetic distance (Mantel’s *r* = .22, *p* = .00001), demonstrating that geographical distance represents a barrier to gene flow for *M. germanica.*

**Table S1.** Characteristics of 20 nuclear microsatellite loci 67 *M.germanica* populations, 2212 individuals.

| Locus | Number of alleles | Frequency of null alleles | Ho | Hs | F_IS_ | F_IT_ | F_ST_ |
| --- | --- | --- | --- | --- | --- | --- | --- |
| Mg451 | 14 | 0.04 | 0.1954 | 0.3501 | 0.4419 | 0.5083 | 0.5049 |
| Mg452 | 7 | 0.043 | 0.2044 | 0.2771 | 0.2621 | 0.6208 | 0.6175 |
| Mg457 | 3 | 0.027 | 0.0738 | 0.1419 | 0.4798 | 0.6311 | 0.6278 |
| Mg459 | 9 | 0.028 | 0.1731 | 0.2572 | 0.3269 | 0.5186 | 0.5152 |
| Mg480 | 15 | 0.041 | 0.1681 | 0.2902 | 0.4208 | 0.5568 | 0.5533 |
| Mg488 | 8 | 0.036 | 0.2583 | 0.3974 | 0.3501 | 0.4469 | 0.4434 |
| Mg489 | 7 | 0.031 | 0.0432 | 0.0699 | 0.3818 | 0.8161 | 0.8140 |
| Mg493 | 22 | 0.17 | 0.2248 | 0.3573 | 0.3707 | 0.5694 | 0.5659 |
| Mg495 | 4 | 0.036 | 0.0721 | 0.1151 | 0.3738 | 0.7529 | 0.7503 |
| Mg498 | 6 | 0.038 | 0.1183 | 0.2000 | 0.4086 | 0.6392 | 0.6359 |
| Mg499 | 5 | 0.023 | 0.1974 | 0.3029 | 0.3483 | 0.3646 | 0.3614 |
| Mg500 | 11 | 0.038 | 0.1237 | 0.2143 | 0.4230 | 0.6308 | 0.6276 |
| Mg549 | 5 | 0.037 | 0.1862 | 0.2505 | 0.2568 | 0.6009 | 0.5975 |
| Mg442 | 7 | 0.044 | 0.1786 | 0.2675 | 0.3324 | 0.6399 | 0.6367 |
| Mg444 | 10 | 0.044 | 0.0672 | 0.1383 | 0.5140 | 0.7551 | 0.7525 |
| Mg455 | 7 | 0.037 | 0.2054 | 0.3115 | 0.3408 | 0.5291 | 0.5256 |
| Mg462 | 15 | 0.19 | 0.1352 | 0.2291 | 0.4097 | 0.6016 | 0.5983 |
| Mg502 | 10 | 0.040 | 0.1573 | 0.2144 | 0.2663 | 0.4768 | 0.4733 |
| Mg504 | 6 | 0.036 | 0.1754 | 0.2331 | 0.2473 | 0.6323 | 0.6291 |
| Mg507 | 8 | 0.038 | 0.1374 | 0.1894 | 0.2744 | 0.6627 | 0.6596 |

**Table S2.** Characteristics of six chloroplast microsatellite loci for 67 *M.germanica* populations, 2212 individuals. Ho, observed heterozygosity, the Shannon-Wiener Diversity index; Hd, Haplotype diversity; *F*_ST_, fixation index of subpopulation to total.

| Locus | Number of alleles | Hd | *F*_ST_ |
| --- | --- | --- | --- |
| psbA | 9 | 0.6323 | 0.1495 |
| psbC | 6 | 0.6408 | 0.1514 |
| trnC1 | 2 | 0.5721 | 0.1100 |
| trnC2 | 2 | 0.6128 | 0.1455 |
| trnH | 7 | 0.6301 | 0.1568 |
| trnL | 4 | 0.5838 | 0.1053 |

**Table S3.** Global analysis of molecular variance (AMOVA) with 20 nuclear loci, 67 populations and 2212 individuals of *M. germanica.* Df (degrees of freedom), SS (Sum of squares), MS (Mean square).

| **Source** | **Df** | **SS** | **MS** | **Sigma** | **% covariance** |
| --- | --- | --- | --- | --- | --- |
| **Among Pops** | 66 | 6415.960 | 267.332 | 3.018 | 49 |
| **Within Pops** | 2212 | 7009.743 | 3.095 | 3.095 | 51 |
| **Total** | 2236 | 13425.702 |  | 6.113 | 100 |

**Table S4.** Global analysis of molecular variance (AMOVA) with 6 chloroplast loci, 67 populations and 2212 individuals of *M. germanica.* Df (degrees of freedom), SS (Sum of squares), MS (Mean square).

| **Source** | **Df** | **SS** | **MS** | **Est.Var** | **%** |
| --- | --- | --- | --- | --- | --- |
| **Among Pops** | 66 | 3782.940 | 54.8252114 | 1.7166617 | 75.25 |
| **Within Pops** | 2212 | 1212.574 | 0.5645129 | 0.5645129 | 24.75 |
| **Total** | 2236 | 4995.513 |  | 2.2811747 | 100 |

**Table S5**. Pairwise relative migration probabilities among river catchments estimated from nuclear microsatellites using MIGRATE-n

|  | Po | Isar | Oder | Lech | Inn | Drau | Aare | Rhine | Rhone | Adige | Tagliamento | Danube |
| --- | --- | --- | --- | --- | --- | --- | --- | --- | --- | --- | --- | --- |
| Po | — | 0,54 | 0,54 | 0,62 | 0,38 | 0,63 | 0,41 | 0,39 | 0,26 | 0,28 | 0,71 | 0,35 |
| Isar | 0,61 | — | 0,26 | 0,43 | 0,36 | 0,47 | 0,40 | 0,30 | 0,32 | 0,34 | 0,44 | 0,39 |
| Oder | 0,43 | 0,35 | — | 0,65 | 0,40 | 0,64 | 0,51 | 0,27 | 0,42 | 0,26 | 0,50 | 0,75 |
| Lech | 0,50 | 0,35 | 0,45 | — | 0,50 | 0,44 | 0,43 | 0,69 | 0,56 | 0,37 | 0,44 | 0,37 |
| Inn | 0,37 | 0,30 | 0,56 | 0,69 | — | 0,27 | 0,29 | 0,41 | 0,32 | 0,63 | 0,38 | 0,28 |
| Drau | 0,57 | 0,46 | 0,27 | 0,52 | 0,32 | — | 0,36 | 0,52 | 0,60 | 0,73 | 0,40 | 0,32 |
| Aare | 0,55 | 0,37 | 0,26 | 0,47 | 0,33 | 0,29 | — | 0,40 | 0,26 | 0,42 | 0,42 | 0,50 |
| Rhine | 0,62 | 0,35 | 0,59 | 0,35 | 0,25 | 0,81 | 0,40 | — | 0,29 | 0,22 | 0,66 | 0,52 |
| Rhone | 0,34 | 0,49 | 0,46 | 0,42 | 0,84 | 0,45 | 0,68 | 0,23 | — | 0,50 | 0,59 | 0,68 |
| Adige | 0,57 | 0,26 | 0,37 | 0,39 | 0,65 | 0,26 | 0,28 | 0,41 | 0,54 | — | 0,40 | 0,65 |
| Tagliamento | 0,55 | 0,63 | 0,51 | 0,72 | 0,26 | 1,00 | 0,60 | 0,42 | 0,51 | 0,34 | — | 0,32 |
| Danube | 0,45 | 0,28 | 0,52 | 0,50 | 0,31 | 0,32 | 0,55 | 0,46 | 0,61 | 0,47 | 0,19 | — |

Table S6. Pairwise relative migration probabilities among river catchments estimated from chloroplast microsatellites using MIGRATE-n

|  | Po | Isar | Oder | Lech | Inn | Drau | Aare | Rhine | Rhone | Adige | Tagliamento | Danube |
| --- | --- | --- | --- | --- | --- | --- | --- | --- | --- | --- | --- | --- |
| Po | — | 0,004 | 0,007 | 0,006 | 0,011 | 0,007 | 0,008 | 0,005 | 0,005 | 0,014 | 0,007 | 0,007 |
| Isar | 0,013 | — | 0,008 | 0,005 | 0,008 | 0,003 | 0,004 | 0,004 | 0,005 | 0,010 | 0,006 | 0,008 |
| Oder | 0,006 | 0,005 | — | 0,005 | 0,006 | 0,011 | 0,006 | 0,010 | 0,012 | 0,008 | 0,007 | 0,007 |
| Lech | 0,007 | 0,006 | 0,004 | — | 0,015 | 0,007 | 0,005 | 0,008 | 0,009 | 0,007 | 0,007 | 0,008 |
| Inn | 0,014 | 0,008 | 0,006 | 0,010 | — | 0,011 | 0,006 | 0,011 | 0,008 | 0,012 | 0,010 | 0,012 |
| Drau | 0,006 | 0,007 | 0,006 | 0,010 | 0,006 | — | 0,005 | 0,008 | 0,004 | 0,008 | 0,008 | 0,010 |
| Aare | 0,005 | 0,005 | 0,006 | 0,003 | 0,005 | 0,015 | — | 0,008 | 0,006 | 0,007 | 0,013 | 0,007 |
| Rhine | 0,006 | 0,014 | 0,006 | 0,005 | 0,015 | 0,007 | 0,007 | — | 0,007 | 0,007 | 0,007 | 0,006 |
| Rhone | 0,008 | 0,006 | 0,005 | 0,005 | 0,004 | 0,008 | 0,007 | 0,006 | — | 0,009 | 0,010 | 0,012 |
| Adige | 0,004 | 0,009 | 0,006 | 0,010 | 0,005 | 0,011 | 0,007 | 0,011 | 0,006 | — | 0,004 | 0,008 |
| Tagliamento | 0,012 | 0,004 | 0,006 | 0,008 | 0,005 | 0,008 | 0,007 | 0,013 | 0,012 | 0,009 | — | 0,009 |
| Danube | 0,005 | 0,009 | 0,003 | 0,009 | 0,008 | 0,005 | 0,005 | 0,005 | 0,011 | 0,006 | 0,008 | — |

Table S7. Pairwise relative migration probabilities among river catchments estimated from nuclear microsatellites using BA3.

|  | Po | Isar | Oder | Lech | Inn | Drau | Aare | Rhine | Rhone | Adige | Tagliamento | Danube |
| --- | --- | --- | --- | --- | --- | --- | --- | --- | --- | --- | --- | --- |
| Po | 0,688 | 0,012 | 0,033 | 0,013 | 0,011 | 0,036 | 0,027 | 0,013 | 0,024 | 0,031 | 0,008 | 0,038 |
| Isar | 0,034 | 0,730 | 0,038 | 0,021 | 0,016 | 0,039 | 0,043 | 0,024 | 0,041 | 0,042 | 0,016 | 0,036 |
| Oder | 0,001 | 0,001 | 0,674 | 0,001 | 0,010 | 0,006 | 0,004 | 0,001 | 0,003 | 0,006 | 0,001 | 0,008 |
| Lech | 0,047 | 0,023 | 0,041 | 0,758 | 0,020 | 0,042 | 0,038 | 0,030 | 0,042 | 0,038 | 0,018 | 0,039 |
| Inn | 0,069 | 0,043 | 0,041 | 0,021 | 0,779 | 0,047 | 0,047 | 0,025 | 0,046 | 0,049 | 0,033 | 0,046 |
| Drau | 0,001 | 0,001 | 0,008 | 0,001 | 0,001 | 0,673 | 0,004 | 0,001 | 0,003 | 0,006 | 0,000 | 0,008 |
| Aare | 0,013 | 0,006 | 0,023 | 0,006 | 0,005 | 0,022 | 0,694 | 0,005 | 0,013 | 0,022 | 0,004 | 0,026 |
| Rhine | 0,080 | 0,035 | 0,045 | 0,036 | 0,045 | 0,044 | 0,041 | 0,844 | 0,056 | 0,045 | 0,015 | 0,043 |
| Rhone | 0,016 | 0,008 | 0,032 | 0,009 | 0,008 | 0,027 | 0,023 | 0,008 | 0,699 | 0,028 | 0,008 | 0,027 |
| Adige | 0,001 | 0,001 | 0,008 | 0,001 | 0,001 | 0,007 | 0,004 | 0,001 | 0,003 | 0,673 | 0,001 | 0,008 |
| Tagliamento | 0,049 | 0,139 | 0,051 | 0,133 | 0,113 | 0,050 | 0,072 | 0,048 | 0,070 | 0,055 | 0,895 | 0,046 |
| Danube | 0,001 | 0,001 | 0,008 | 0,001 | 0,001 | 0,007 | 0,004 | 0,001 | 0,003 | 0,006 | 0,001 | 0,675 |
